# The abscopal effect of IRE combined with anti–PD-1 achieves local ablation and systemic control of PDAC

**DOI:** 10.64898/2026.05.04.722535

**Authors:** Qizhen Cao, Zhenzhen Xun, Yitao Tang, Jiakai Hou, Boping Jing, Ping Pan, Jing Zhang, Shiaw-Yih Lin, Sanjay Gupta, Jared K. Burks, Huamin Wang, James P. Long, Han Liang, Weiyi Peng, Chun Li

**Author notes:** Correspondence: Chun Li, PhD. Authors’ Disclosures. No conflict of interest to declare.

## Abstract

Irreversible electroporation (IRE) has shown promise for treating pancreatic ductal adenocarcinoma (PDAC), but whether IRE can induce an abscopal effect is not established. We demonstrated that the combination of IRE and anti–PD-1 antibody could trigger robust abscopal effects in preclinical models of metastatic PDAC. Data from multiple *in vivo* models, RNA-seq, scRNA-seq, and spatial immunofluorescence provide compelling evidence that IRE induced mitochondrial dysfunction and cellular stress, which triggered activation of the cGAS-STING pathway and subsequent systemic antitumor effects. IRE also led to inflammatory response characterized by tumor infiltration of myeloid cells and their polarization toward M1 state, turning immunologically “cold” tumors into “hot” tumors. Moreover, the presence of T cell/B cell clusters in tumors from mice treated with IRE plus αPD-1 and the lack of antitumor efficacy in B cell knockout mice bearing orthotopic murine PDAC tumors indicate that B cells play an important role in IRE-mediated systemic antitumor immunity.

**Significance:** This study shows that IRE plus a checkpoint inhibitor represents a promising therapeutic strategy for PDAC and supports advancing this treatment toward clinical translation. Our data also support potential combination strategies with immunomodulatory agents that can recruit and reprogram B cells to support T cell activation and cytotoxic effector functions.

## INTRODUCTION

The abscopal effect refers to the phenomenon in which localized cancer therapies, such as radiotherapy or local ablation techniques, not only shrink the primary tumor but also elicit regression of distant, untreated metastases. Initially described by Mole in 1953 (1), the abscopal effect has long been considered rare and unpredictable, with most studies focused on radiotherapy (2,3). Recently, interventional techniques such as photothermal ablation (4,5) and irreversible electroporation (IRE) (6) have attracted increasing attention due to their ability to provide minimally invasive solution for *in situ* vaccination.

Evidence has highlighted the pivotal role of the immune system in mediating the abscopal effect (7,8). Local ablation therapy induces immunogenic cell death and releases tumor-associated antigens and damage-associated molecular patterns (DAMPs) that activate antigen-presenting cells. Activated antigen-presenting cells then prime tumor-reactive cytotoxic T cells, which can migrate and attack distant tumor cells (9). In addition, induction of DAMPs, such as cytoplasmic double-stranded DNAs (dsDNAs), can trigger activation of cyclic GMP-AMP synthase (cGAS)–stimulator of interferon genes (STING) signaling, enhancing interferon responses that support T cell activation (10). Moreover, local ablation therapy can modulate the local tumor microenvironment (TME) by promoting tumor infiltration of immune cells such as neutrophils and dendritic cells (11). Preclinical and clinical data increasingly support the use of immune checkpoint inhibitors alongside local ablation therapy to increase the rate of abscopal response compared to local ablation therapy alone (12,13).

IRE is a novel form of local tumor ablation that uses short high-voltage electric pulses to induce tumor cell death by disrupting cell membranes (14). Unlike thermal ablation methods, IRE preserves surrounding critical tissues, making it particularly advantageous for treating tumors located in sensitive anatomical regions such as the liver, pancreas, and prostate (15,16). IRE was first introduced in the clinic in 2011 (17). Martin et al. (18) reported a study of 200 patients with locally advanced pancreatic ductal adenocarcinoma (PDAC) and showed longer survival with surgery plus IRE than with IRE alone. However, patients in both treatment groups commonly experienced recurrence with metastatic disease (18). Currently, there are multiple active clinical trials registered in the ClinicalTrials.gov database involving IRE alone or combined with chemotherapy or radiotherapy for PDAC.

PDAC is a highly aggressive malignancy with a strong tendency to metastasize, most commonly to the liver, and metastatic PDAC is associated with a dismal 5-year survival rate of approximately 3% (19,20). While IRE as a local ablation therapy has been shown to induce immunologic cell death and promote systemic antitumor immunity (6,16), IRE’s potential to induce an abscopal effect to control metastatic spread remains understudied. PDAC is an immunologically “cold” tumor with limited responses to immune checkpoint inhibitors (21,22). Our previous studies have demonstrated that combining IRE with immune checkpoint inhibitors such as anti–PD-1 antibody (αPD-1) enhances CD8⁺ T cell infiltration and significantly prolongs survival in orthotopic murine models of PDAC (6). However, the ability of IRE combined with αPD-1 (IRE + αPD-1) to induce regression of distant metastases has not been thoroughly investigated.

In this study, we demonstrated that IRE + αPD-1 induced strong abscopal effect in multiple model systems. We used unbiased approaches, including RNA sequencing (RNA-seq) and single-cell RNA-seq (scRNA-seq), to gain insights into the possible mechanisms of action of the observed abscopal effects. We performed a spatial proteomics study with sequential immunofluorescence (SeqIF) staining, which provided evidence supporting multiple roles of IRE in stroma modulation. These roles included cGAS-STING pathway activation, enhancement of interferon response, and induction of an immunologically “hot” tumor characterized by infiltration of abundant immune cells in the TME. In particular, our findings highlight the essential role of B cells in maintaining adaptive immunity. Our data suggest potential novel combination strategies for enhancing abscopal effects against PDAC with metastatic disease.

## RESULTS

### IRE + αPD-1 Induced Robust Abscopal Effects Against Distant KRAS* Tumors Inoculated Ectopically or Orthotopically

To study abscopal effects, we first inoculated KRAS* cells subcutaneously into the right flank and left flank at the same time in female C57BL/6 mice. When tumors in the right flank reached approximately 7 to 8 mm in longest diameter, they were treated with IRE, while tumors in the left flank were left untreated. For immunotherapy, mice were injected with αPD-1 intraperitoneally at a dose of 100 µg/mouse, 3 times a week for 2 weeks. Mice not treated with IRE or not treated with αPD-1 were used as controls (**Fig. 1A**). The mice in the IRE + αPD-1 group had a median survival more than twice as long as the median survival with each monotherapy (80 days with combination vs. 31 days with IRE alone and 33 days with αPD-1 alone). IRE or αPD-1 monotherapy only moderately prolonged the median survival compared to the median survival in the untreated control group (24 days). Significantly, all mice in the monotherapy groups died by day 60, whereas IRE + αPD-1 induced complete response in three of seven (43%) treated mice (**Fig. 1B**). Individual tumor growth curves showed that three of seven (43%) abscopal tumors in the IRE + αPD-1 group had a complete response by 70 days after initiation of IRE treatment. In comparison, neither IRE monotherapy nor αPD-1 monotherapy resulted in eradication of abscopal tumors (**Fig. 1C**). Similar trend was observed for the IRE tumors: the rates of complete response were 71% (five of seven) in the IRE + αPD-1 group, 43% (three of seven) in the IRE alone group, and 0% for the no treatment control and the αPD-1 group alone groups (**Fig. S1**). These data indicate that long-term survival was primarily a result of eradication of tumor at distant sites and that αPD-1 was necessary but not sufficient for effective local tumor control and a robust abscopal effect.

**Figure 1.**
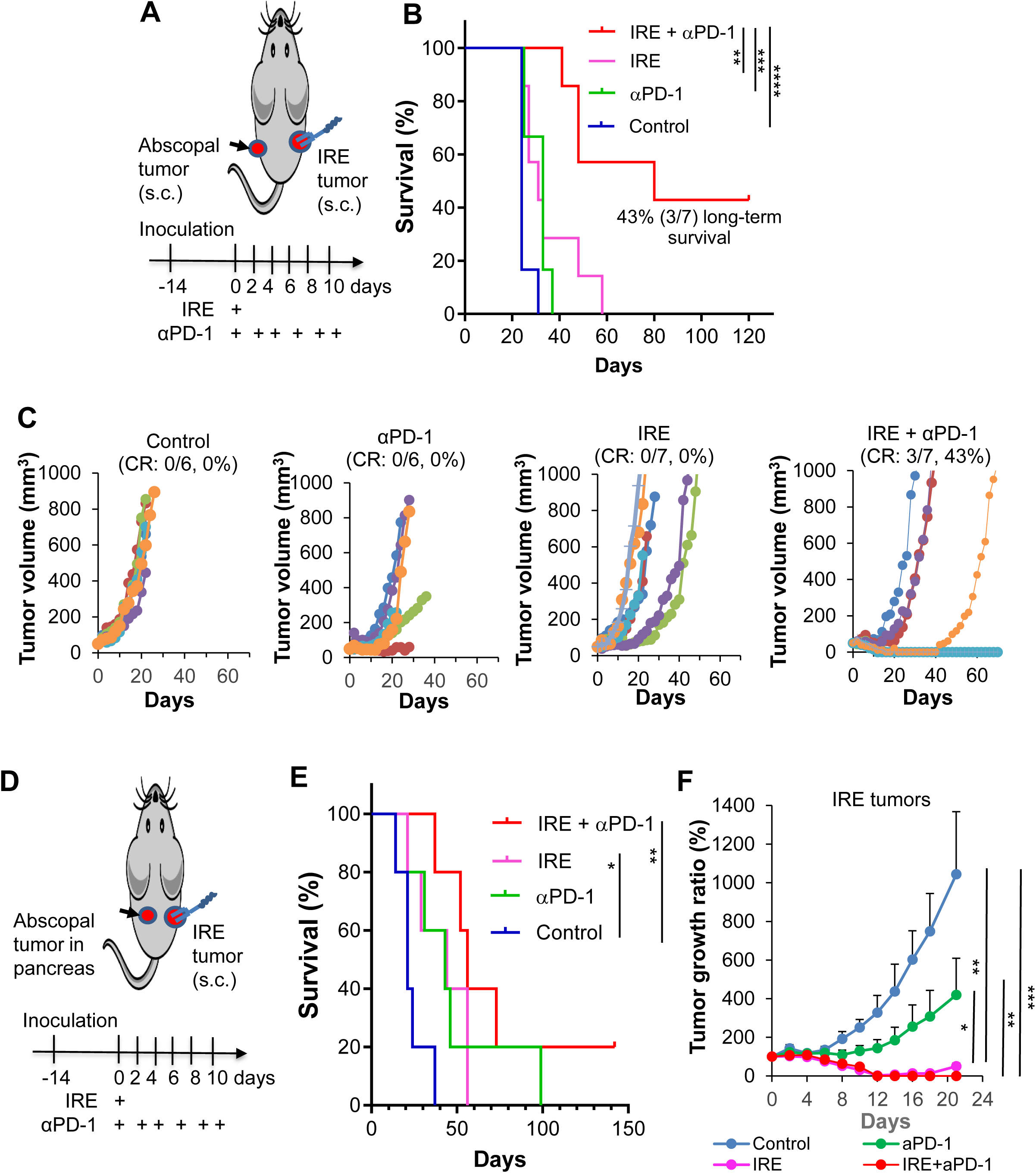
IRE + αPD-1 induced robust abscopal effects in KRAS* tumor–bearing mice. (**A-C**) Abscopal effect in ectopic tumors. (**A**) Schematic diagram illustrates the experimental setup in female C57BL/6 mice with bilateral KRAS* PDAC tumor cells implanted subcutaneously in both flanks. IRE was applied to tumors in the right flank. (**B**) Kaplan-Meier survival curves (n=6-7, log-rank test, *p<0.05, **p<0.01, ***p<0.001, ****p<0.0001). (**C**) Tumor growth curves for the contralateral abscopal tumors (n=6-7). CR, complete response. (**D-F**) Abscopal effect in orthotopic tumors. (**D**) Schematic diagram illustrates the experimental setup in female C57BL/6 mice with KRAS* PDAC tumor cells implanted subcutaneously in the right flank and orthotopically in the pancreas. IRE was applied to tumors in the right flank. (**E**) Kaplan-Meier survival curves (n=5, log-rank test, *p<0.05, **p<0.01). (**F**) Tumor growth curves for the IRE tumors (n=5). (**A-F**) In both experiments, IRE was instituted with the following parameters: voltage, 1200 V; pulse duration, 100 µs; pulse repetition frequency, 1 Hz; number of pulses, 99. αPD-1 was injected intraperitoneally at 100 µg/mouse/injection every other day for a total of 6 doses.

Next, we inoculated KRAS* cells subcutaneously into the right flank and orthotopically into the pancreas in female C57BL/6 mice. When tumors in the right flank reached approximately 7 to 8 mm in longest diameter, they were treated with IRE in conjunction with six doses of αPD-1 administered intraperitoneally. Pancreatic tumors were left untreated. Mice not treated with IRE and not treated with αPD-1 were used as controls (**Fig. 1D)**. The median survival times for untreated control, IRE alone, αPD-1 alone, and IRE + αPD-1 were 21 days, 44 days, 43 days, and 56 days, respectively (**Fig. 1E**). IRE alone and IRE + αPD-1 significantly increased survival compared to no treatment (log-rank test, p<0.05, p<0.01). The combination treatment group not only had longer median survival than each monotherapy group but also included a mouse (1/5, 20%) with a durable response (**Fig. 1E**). Both IRE-based treatments successfully delayed growth of the IRE tumors compared to no treatment or αPD-1 alone (**Fig. 1F**). These data indicate that IRE + αPD-1 displayed antitumor activity against orthotopically inoculated abscopal tumors and indicate that while IRE effectively eliminated local tumors, survival was primarily governed by the robustness of the abscopal effects.

### *In Vitro* RNA-seq Analysis Revealed Metabolic Reprogramming and Activation of Pro-inflammatory Pathways

To gain insights into the impact of IRE on tumor cells, we used RNA-seq analysis to characterize transcriptomic changes in KRAS* cells upon IRE treatment. **Figures S2A and S2B** show the results of principal component analysis of cells collected at 4 h and 24 h after IRE at 400 V with 10 pulses (10p IRE) or 20 pulses (20p IRE) versus untreated control cells.

Unsupervised cluster analysis showed that both 10p IRE and 20p IRE treatments induced gene expression changes in KRAS* cells, distinguishing them from the control group. Moreover, the principal component analysis plots showed that within-group samples were clustered together, validating the quality and reliability of our data. The numbers of differentially expressed genes (DEGs) (|log_2_FC| > 2.0 & false discovery rate < 0.01) at 4 h after IRE for 10p IRE versus untreated control, 20p IRE versus untreated control, and 20p IRE versus 10p IRE are presented in a Venn diagram (**Fig. S2C)**. A significant overlap exists between the genomic responses to 10p IRE and 20p IRE. Specifically, 34.8% (243/698) of upregulated genes and 20.3% (50/246) of downregulated genes are shared, indicating similarity in cellular responses regardless of the number of pulses.

Volcano maps were constructed to visualize the distribution of DEGs for 10p IRE versus control and 20p IRE versus control (**Fig. 2A**). The top 21 protein-encoding DEGs that were upregulated at 4 h after both 10p IRE and 20p IRE are listed in **Table S1**. Proteins encoded by these 21 genes can be broadly classified into four categories: proteins involved in antigen presentation (e.g., *Mmp25, Hspa1a*), stress response (e.g., *Npas4, Aldh3b2, Matn4, Mc1r, Egr2,* and *Prkcg*), immune response/recruitment of immune cells/inflammation (e.g., *Pecam1, Ampd1, Itgam, Slc8a2/3, Cxcl2, Aoc3, Pde4b,* and *Crabp2*), and stroma remodeling (e.g., *Pecam1*, *Gdf9*, and *Edn1*). Of note, *Itgam* (*CD11b*), which is a marker for myeloid cells that are abundant in the TME of PDAC (23), was upregulated in IRE-treated PDAC cells. *Cxcl2*, which acts as a powerful chemoattractant for rapid recruitment of neutrophils, was also upregulated. Increased *CD11b* and *Cxcl2* expression suggest that after IRE, tumor cell injury may trigger rapid recruitment of neutrophils and/or macrophages to initiate acute inflammation and innate immune response. Among the upregulated genes, only *Hspa1α* was upregulated under all four treatment conditions (10p IRE, 4 h; 20p IRE, 4 h; 10p IRE, 24 h; and 20p IRE, 24 h). *Hspa1α* encodes the highly inducible 70-kDa heat shock protein (Hsp70).

**Figure 2.**
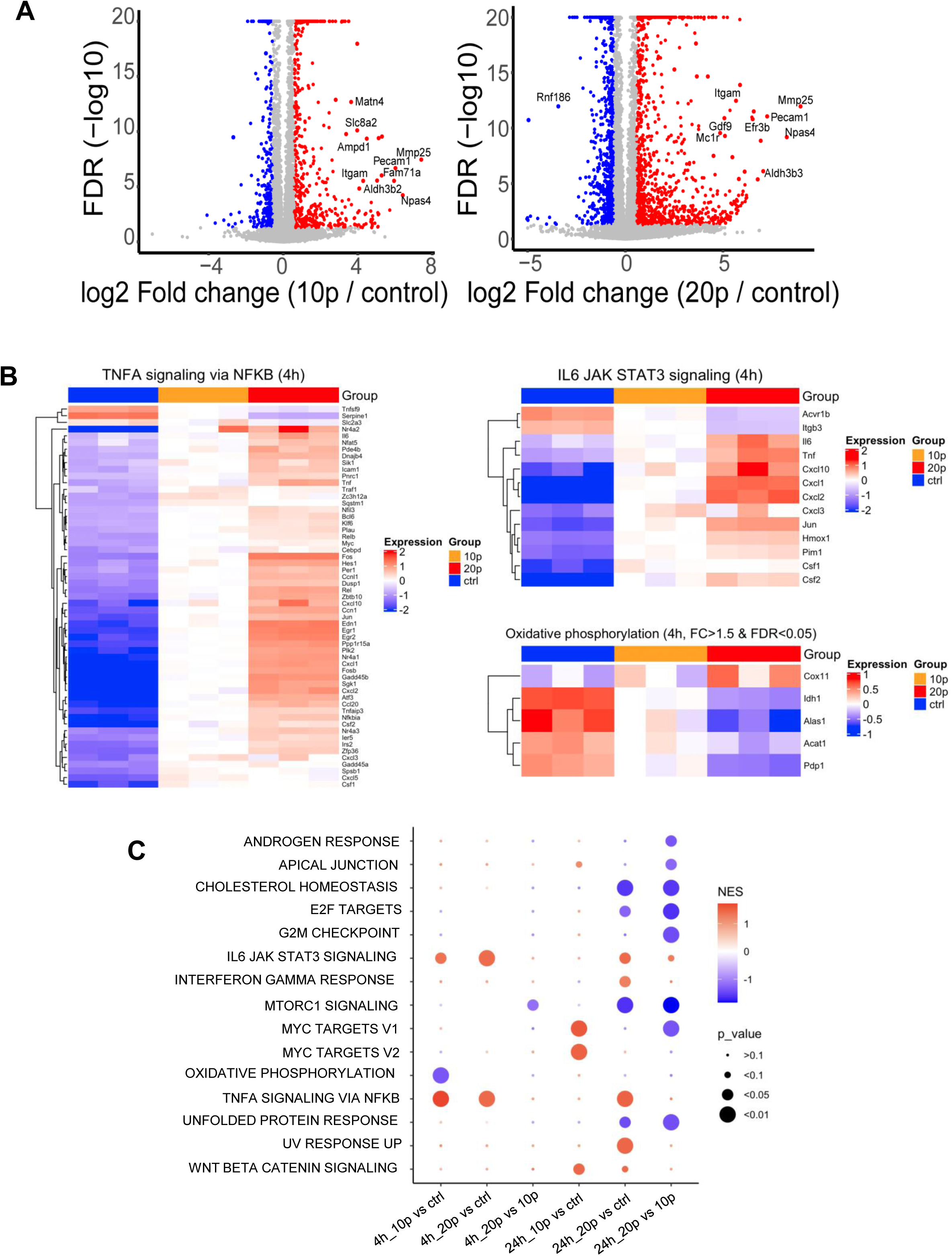
RNA-seq analysis of KRAS* cells at 4 h after IRE (400 V). (**A**) Volcano plot showing DEGs with significantly decreased (blue) or increased (red) expression (p≤0.05). FDR, false discovery rate. (**B**) Heatmap illustrating key signaling pathways identified by GSEA. (**C**) Results of GO enrichment analysis based on biological processes.

Gene set enrichment analysis (GSEA) revealed that DEGs identified in the IRE groups were significantly enriched in three signaling pathways: TNF-α via NF-κB, IL-6/JAK/STAT3, and oxidative phosphorylation (OXPHOS) (**Fig. 2B)**. A scatterplot generated from Gene Ontology (GO) enrichment analysis is presented in **Figure 2C**. Again, the TNF-α via NF-κB pathway and IL-6/JAK/STAT3 pathway were upregulated at 4 h after 10p IRE and 20p IRE and at 24 h after 20p IRE. OXPHOS signaling was downregulated at 4 h after 10p IRE. The GO enrichment analysis also revealed upregulation of several signaling pathways that stimulate cell proliferation (i.e., WNT beta catenin signaling at 24 h after 10p IRE and 20p IRE, Myc targets V1/V2 at 24 h after 10p IRE). These data indicate that it is critical for IRE to completely eliminate the primary tumor because residual tumor cells left behind may recur at a faster growth rate due to changes in signaling pathways.

### IRE Decreased the Metabolic Activity of Cancer Cells by Suppressing Mitochondrial Respiration and Glycolysis

Given that both GSEA and GO enrichment analysis indicated an impact of IRE on OXPHOS activity, we assessed the influence of IRE on the metabolic activity of cancer cells. At 4 h after IRE (400 V, 10p or 20p), the oxygen consumption rate (**Fig. 3A**) and extracellular acidification rate (**Fig. 3B**) of KRAS* PDACs cells were significantly lower than the rates in untreated control cells. Oxygen consumption rate reflects mitochondrial respiration via OXPHOS, while extracellular acidification rate indicates glycolysis activity. Thus, IRE significantly suppressed mitochondrial respiration and glycolytic activity of cancer cells. Specific metabolic parameters, including ATP production (**Fig. 3C**), were significantly lower in IRE-treated KRAS* cells than in untreated control cells. IRE-treated KRAS* cells had a significantly lower basal respiration rate, maximal respiration rate, and spare respiratory capacity (mitochondrial reserve capacity for producing ATP) than control cells in a pulse number-dependent manner (**Figs. S3A-S3C**). Taken together, these findings indicated that IRE with a number of pulses as low as 10 caused mitochondrial dysfunction of PDAC cells.

**Figure 3.**
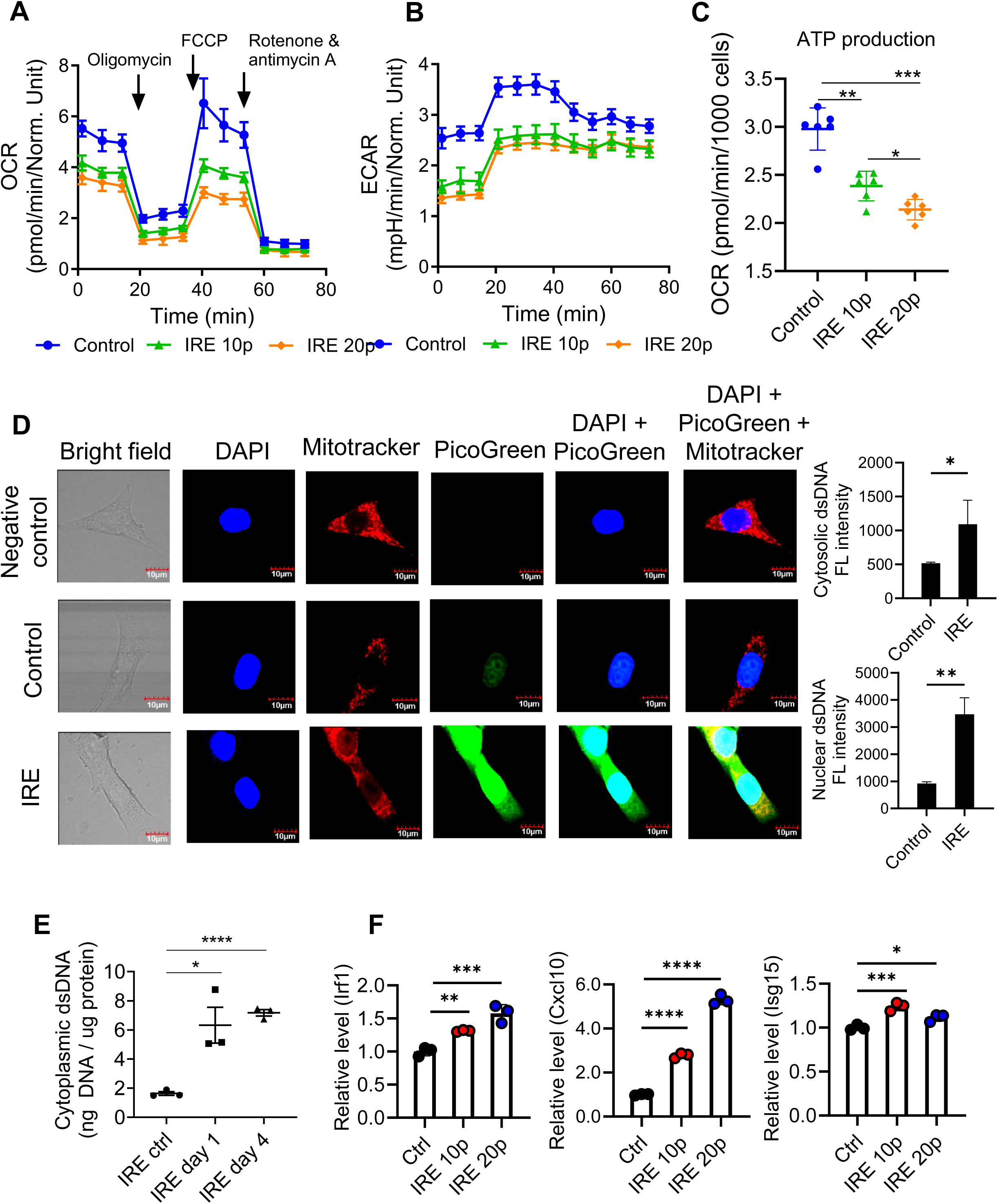
IRE induced mitochondrial dysfunction and induced cytosolic release of dsDNA to activate the cGAS-STING sensing pathway. (**A**) Mitochondrial oxygen consumption rate (OCR) and (**B**) extracellular acidification rate (ECAR) in KRAS* cells 4 h after 10p IRE or 20p IRE (400 V). (**C**) Effect of IRE on mitochondrial ATP production. Data are normalized to cell number at the end of the assay (pmol/min/1000 cells). Data are expressed as mean ± SD (n=6/group). *p<0.05, **p<0.01, ***p<0.001 (one-way ANOVA). (**D**) Representative confocal fluorescence microscopic images of cytoplasmic dsDNA and corresponding quantification of fluorescence (FL) signal intensity in cytosol and nuclei. PANC-1 cells were stained with PicoGreen for dsDNA 24 h after IRE (400 V, 20 pulses). Untreated cells were used as a control. The negative control was cells not stained with PicoGreen. Data are expressed as mean ± SD (n=4). *p<0.05, *p<0.01, (Student’s t test). (**E**) Analysis of dsDNA from IRE-treated KRAS* tumors using the Qubit dsDNA HS assay kits. KRAS* tumor tissues were collected on day 1 and day 4 after IRE (1200 V, 99 pulses) and processed for dsDNA assays. *p<0.05, ****p<0.0001 (Student’s t test). (**F**) Real-time PCR analysis of the interferon-stimulated genes *Irf1*, *Cxcl10*, and *Isg15* downstream of the cGAS-STING pathway. KRAS* cells were collected 4 h after IRE (400 V, 20 pulses). *p<0.05, **p<0.01, ***p<0.001, ****p<0.0001 (one-way ANOVA). Data are expressed as mean ± SD (n=3/group).

### IRE Caused Cytosolic Accumulation of dsDNA and ssDNA and Activation of cGAS-STING Pathway

Mitochondrial dysfunction may be attributed in part to IRE-induced mitochondrial membrane damage, which leads to mitochondrial DNA leakage. We therefore examined release of DNA into cytosol. Confocal microscopic images showed minimal PicoGreen-labeled dsDNA in untreated control cells. However, at 24 h after IRE, fluorescence signal intensity from PicoGreen significantly increased in the cytoplasm of PANC-1 PDAC cells (**Fig. 3D**). Moreover, there was strong PicoGreen signal in the mitochondria and nuclei of IRE-treated tumor cells (**Fig. 3D**). Similarly, fluorescent signal intensity from OliGreen, a ssDNA-specific dye, was higher in the cytoplasm, mitochondria, and nuclei of IRE-treated PDAC cells than those in the untreated cells (**Fig. S4A**). These data demonstrated that IRE potently induced DNA leakage from mitochondria and nuclei to cytoplasm. Quantification of cytosolic DNA from IRE-treated KRAS* tumors further showed that IRE significantly increased cytosolic dsDNA (**Fig. 3E**) and ssDNA (**Fig. S4B**) at 1 day and 4 days after IRE treatment compared to untreated control tumors.

The presence of cytosolic dsDNA could act as a danger signal, activating the immune system through the cGAS–STING cytosolic DNA–sensing pathway (24,25). ssDNA accumulation is often linked to genomic instability, replication stress, and DNA damage (26). Therefore, the observed presence of cytosolic ssDNA linked IRE-induced DNA damage and replication stress to innate immune and inflammatory responses. To determine whether the cGAS–STING pathway was activated, KRAS* cells were treated with 10p IRE and 20p IRE. Four hours after treatment, real-time PCR was used to analyze expression levels of interferon-stimulated genes downstream of the cGAS-STING signaling pathway, including *Irf1*, *Cxcl10*, and *Isg15*. As shown in **Figure 3F**, IRE significantly increased the expression levels of these genes relative to the levels in the corresponding controls. Cxcl10 can act as a powerful signaling chemokine to recruit immune cells like T cells, natural killer (NK) cells, monocytes, and dendritic cells (27). Elevated Cxcl10 also supports involvement of the TNF-α via NF-κB pathway and the IL-6/JAK/STAT3 pathway, as revealed by the RNA-seq data (**Fig. 2**). These data suggest that immune activation was likely conferred from downstream ISG expression.

### IRE + αPD-1 Induced Abscopal Effects Against Distant Liver Tumor in an Orthotopic KRAS* Tumor Model

PDAC has a high potential for liver metastasis, which is a major contributor to poor prognosis for PDAC patients (28). We therefore developed a liver metastasis model to investigate the impact of local IRE treatment on distant liver tumor to mimic the clinical situation. KRAS* cells were inoculated in both the pancreas and the left lobe of the liver. The primary KRAS* tumor in the pancreas was treated with IRE, while the distant liver tumor was not treated. Six doses of αPD-1 were administered intraperitoneally (**Fig. 4A)**. Mice not treated with IRE or not treated with αPD-1 were used as controls. All mice were euthanized on day 14 after initiation of IRE and/or αPD-1. The volumes of IRE-treated pancreatic tumors in the IRE + αPD-1 group were significantly smaller than those in the untreated control group or αPD-1 alone group (**Fig. 4B).** The mean volumes of liver metastases in the untreated control, αPD-1 alone, IRE alone, and IRE + αPD-1 groups were 111.3±161.6 mm^3^, 39.5±54.3 mm^3^, 53.3±51.6 mm^3^, and 3.6±8.0 mm^3^, respectively, and the metastases in the combination treatment group were tightly clustered with respect to size (**Fig. 4C).** Moreover, four of five (80%) mice in the IRE + αPD-1 group had minimal residual liver metastases (tumor nodule <2 mm^3^), compared with none of five (0%) mice in the untreated control group, two of five (40%) treated with αPD-1 alone, and none of five (0%) treated with IRE alone (**Fig. 4C**). These data indicate that treating primary pancreatic tumors with IRE + αPD-1 could suppress the growth of distant liver metastases.

**Figure 4.**
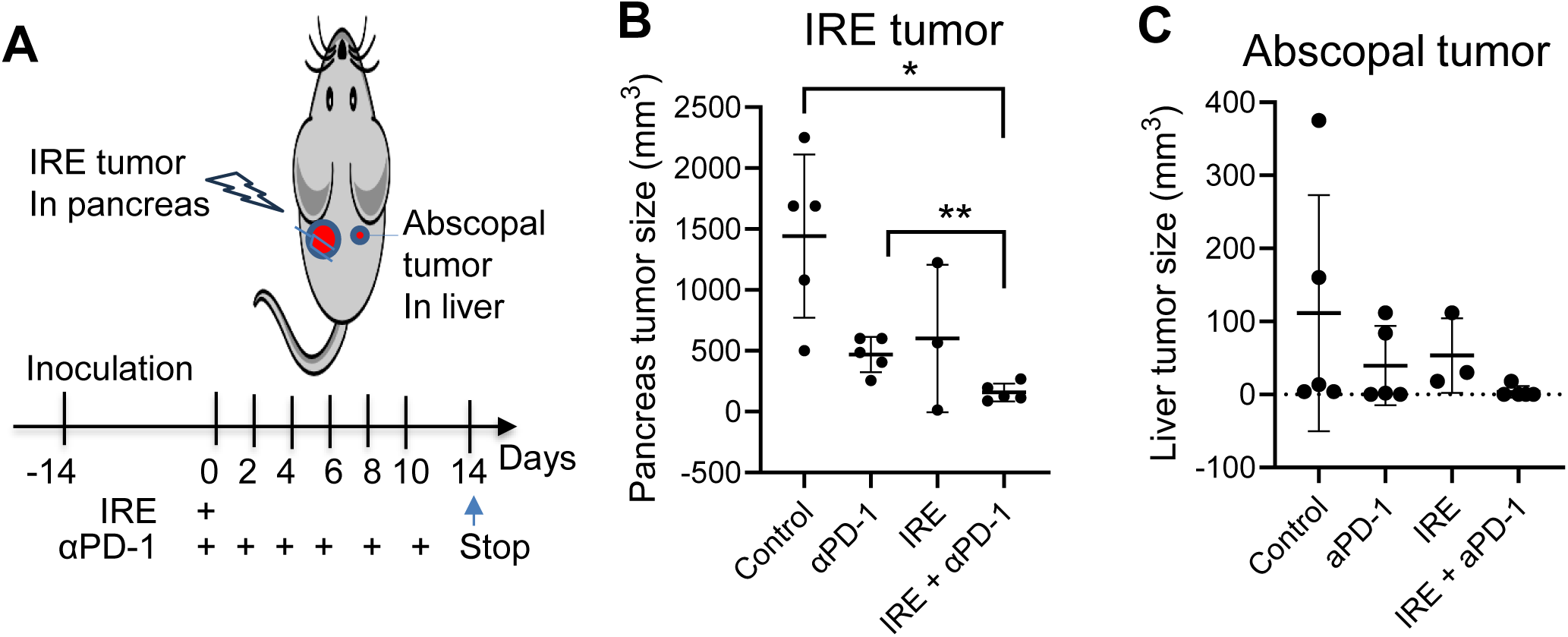
IRE + αPD-1 was effective against distant liver tumor in an orthotopic PDAC model. (**A**) Schematic diagram illustrates the experimental setup in mice with KRAS* tumors in the pancreas and in the liver. IRE treatment (1200 V, 100 µs/pulse, 99 pulses) was applied to the tumors in the pancreas (IRE tumors). αPD-1 (100 µg/mouse/injection) was injected intraperitoneally once every other day for a total of six doses. (**B, C**) Tumor volume measured on day 14, when mice were euthanized, for (**B**) IRE tumors and (**C**) abscopal tumors from the liver. Data are expressed as mean ± SD (n=3-5). *p<0.05, ** p<0.01 (Welch’s t-test).

### IRE + αPD-1 Significantly Impacted the TME of Primary Tumors and Implicated a Crucial Role of B cells in Antitumor Immunity

Our GSEA and GO enrichment analysis data indicated activation of TNF-α signaling via NF-κB (**Figs. 2B** and **2C**), which is intricately linked to inflammation and the recruitment of immune cells (29). We therefore investigated the effects of IRE + αPD-1 on the TME of IRE-treated tumors from the pancreas and abscopal tumors from the liver. We prepared TMAs and performed SeqIF staining using the Lunaphore COMET system. Since IRE treatment leads to necrosis around the tumor ablation zone, tumors were separated into four different zones for analysis: P1, the tumor periphery, farthest from the necrotic zone; P2 and P3, areas between the necrotic zone and the tumor periphery, with P2 closer to the tumor periphery; and P4, the area immediately adjacent to the necrotic zone. An example of an H&E-stained section from an IRE tumor with the zones labeled and a diagram of the zones are presented in **Figure 5A**. Representative H&E-stained TMA slides and corresponding SeqIF images for P4 spots of tumors from each of the four treatment groups are shown in **Figure 5B** and **Figure 5C**, respectively. Higher levels of CD31^+^ tumor vascular vessels and collagen1^+^ collagen were observed in tumors treated with IRE (**Fig. 5C**), suggesting active stromal remodeling. The imaging data clearly reveal that IRE-based treatments were associated with higher tumor infiltration of immune cells, including F4/80^+^ macrophages, CD8^+^/CD4^+^ T cells, and B220^+^ B cells, indicating that IRE converted immunogenically “cold” PDAC tumors to “hot” tumors. Extensive immune infiltration was accompanied by markedly reduced density of PanCK^+^ tumor cells in tumors treated with IRE (**Fig. 5C**). Intriguingly, we observed clusters formed between B220^+^ B cells and CD8^+^ T cells in tumors treated with IRE + αPD-1 (**Fig. 5D**) but not in tumors treated with IRE alone (data not shown). To investigate the role of B cells in IRE + αPD-1 therapy–mediated systemic antitumor immune response, we treated orthotopic KRAS* tumors grown in B cell–knockout transgenic C57BL/6 mice with IRE + αPD-1. The B cell–knockout mice experienced significantly lower median survival than KRAS* tumor–bearing wild type C57BL/6 mice (30) (**Fig. 5E**). These B cell–knockout mice had extensive liver metastases and/or ascites in the intraperitoneal cavity when moribund.

**Figure 5.**
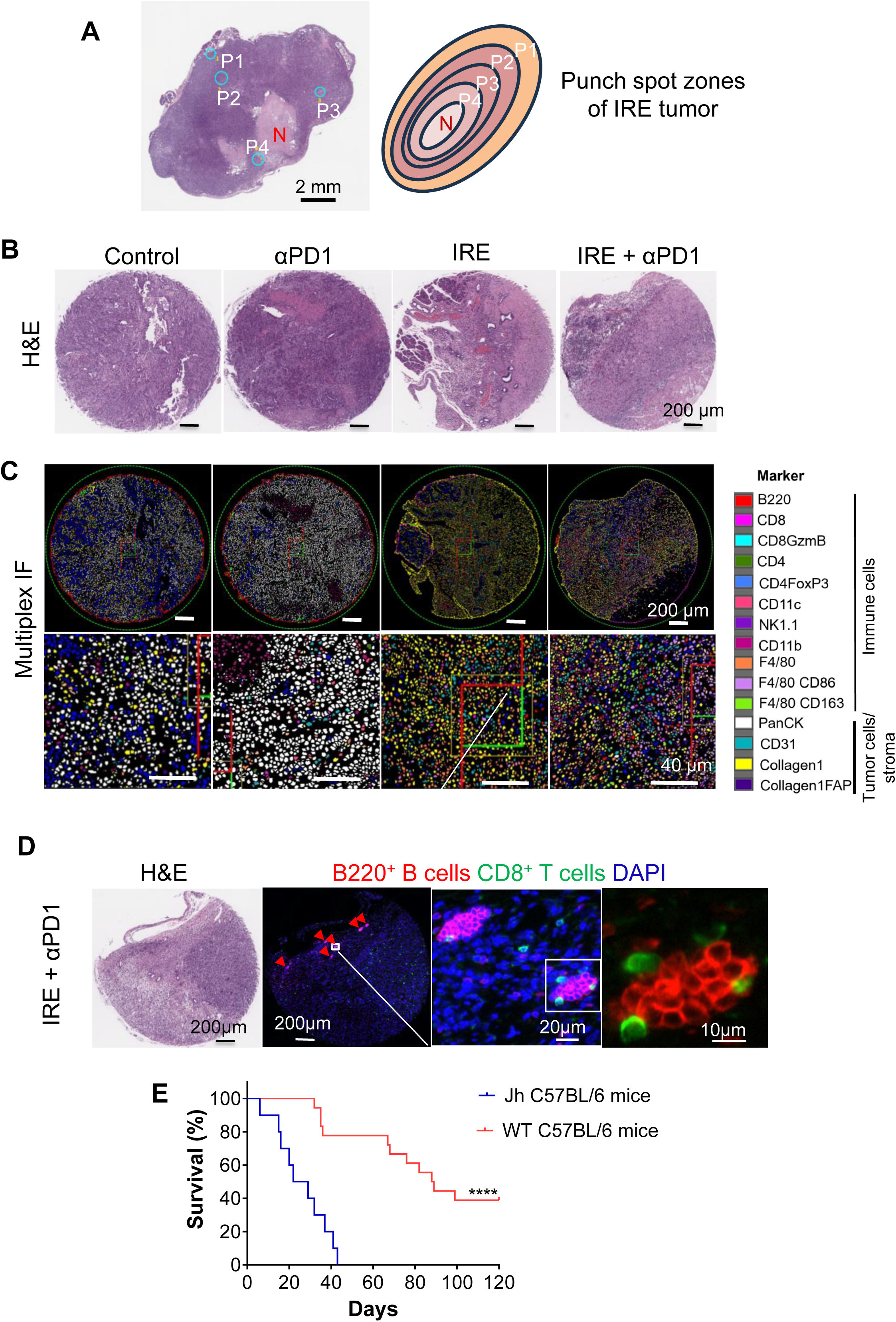
IRE + αPD-1 remodeled the TME to turn immunologically “cold” orthotopic KRAS* PDAC tumors into “hot” tumors. (**A**) Representative photograph of an H&E-stained section from an IRE-treated tumor together with a diagram depicting zones P1-P4. (**B, C**) Representative H&E-stained TMA slides (**B**) and corresponding SeqIF images (**C**) on P4 spots of tumors from each of four treatment groups. (**D**) Representative H&E-stained slides and SeqIF images from zone P4 of an IRE + αPD-1–treated tumor, showing clusters of CD8^+^ T cells and B220^+^ B cells resembling tertiary lymphoid structures. Red arrowheads: clusters of CD8^+^ T cells and B220^+^ B cells at lower magnification. Experimental conditions and setup are outlined in Figure 4A. IRE parameters: voltage, 1200 V; pulse duration, 100 µs; pulse repetition frequency, 1 Hz; number of pulses, 99. (**E**) Kaplan-Meier survival curves of B cell–knockout mice versus wild-type C57BL/6 mice.

Results of a quantitative analysis of different phenotypes of immune cells and stromal cells in zones P3 and P4 are presented as a heatmap in **Figure S5A** and as box and whisker plots in **Figure S5B**. This analysis confirmed the findings of visual inspection. There was a marked increase in the density of immune cells (F4/80^+^ macrophages, B220^+^ B cells, and CD8^+^/CD4^+^T cells) after treatment with IRE + αPD-1. Phenotype analyses showed that the density of F4/80^+^CD86^+^ M1 macrophages in the IRE + αPD-1 group was significantly higher than that in the untreated control, αPD-1 alone, or IRE alone groups (**Fig. S5B**). The density of CD8^+^GzmB^+^ cytotoxic T cells was significantly higher in the IRE + αPD-1 group than in the untreated control and monotherapy groups. In addition, while the density of CD4^+^ cells was higher in the IRE + αPD-1 group than in the monotherapy groups, the density of immunosuppressive CD4^+^FoxP3^+^ regulatory T cells (Tregs) was not affected by IRE-based treatments (**Fig. S5B**).

In addition to higher densities of tumor-associated macrophages (TAMs), B cells, and T cells, tumors treated with IRE + αPD-1 had higher densities of CD31^+^ endothelial cells than the untreated control and αPD-1 alone groups had. The densities of αSMA^+^, αSMA^+^FAP^+^, and collagen1^+^FAP^+^ cancer-associated fibroblasts (CAFs) were significantly higher in the IRE + αPD-1 group than in the αPD-1 alone and/or IRE alone groups, leading to increased collagen 1 deposition in tumors treated with IRE + αPD-1 (**Fig. S5B**). These data indicate that IRE + αPD-1 but not monotherapies actively remodeled tumor stroma and promoted immune cell infiltration and activation.

To confirm the tumor-intrinsic role of IRE-treated tumor cells in recruitment of immune cells, we analyzed macrophage migration in a co-culture system. Macrophage migration was significantly greater with IRE-treated KRAS* cells than with untreated control cells (**Figs. S6A-S6C**). Moreover, co-culture assay showed that RAW264.7 macrophages phagocytosed tumor cells more effectively after murine HY19636 KRAS cells were treated with IRE alone than after these cells were treated with αPD-1 alone (**Fig. S6D**).

Representative H&E-stained slides and corresponding SeqIF images of zones P1 and P2 of the IRE tumors from the pancreas are shown in **Figures S7A and S7B**. Changes in the TME were notable but less extensive than changes in zone P4 (**Fig. 5**). The densities of different immune cells in zones P1 and P2 are shown as box and whisker plots in **Figure S7C**. IRE + αPD-1 significantly increased the density of F4/80^+^CD86^+^ M1 TAMs compared to IRE or αPD-1 monotherapy. IRE + αPD-1 also significantly increased the density of cytotoxic CD8^+^GzmB^+^ T cells compared to IRE monotherapy or untreated control (**Fig. S7C**). Interestingly, the density of immunosuppressive CD4^+^FoxP3^+^ Tregs in zones P1 and P2 was significantly lower in all therapy groups than in the control group (**Fig. S7C**). Similar to what was observed in zones P3 and P4, there was a general trend of higher density of αSMA^+^ CAFs, collagen 1^+^FAP^+^ CAFs, and collagen 1 expression in the IRE + αPD-1 group than in the untreated control or monotherapy groups in zones P1 and P2, which were further away from the IRE needles.

### IRE + αPD-1 Significantly Impacted the TME of Abscopal Liver Metastases

Representative H&E-stained slides and corresponding SeqIF images of liver metastases from each treatment group are presented in **Figures 6A and 6B**. SeqIF images showed higher infiltration of immune cells in the liver metastases in mice treated with IRE + αPD-1, indicating that the combination therapy profoundly impacted the TME of abscopal tumors. Notable findings included higher infiltration of CD11c^+^ dendritic cells, F4/80^+^CD86^+^ M1 TAMs, T cells, B cells at the periphery of tumor nodules and lower density of panCK^+^ tumor cells in tumor nodules in the IRE + αPD-1 group than in the other treatment groups (**Figs. 6A and 6B**).

**Figure 6.**
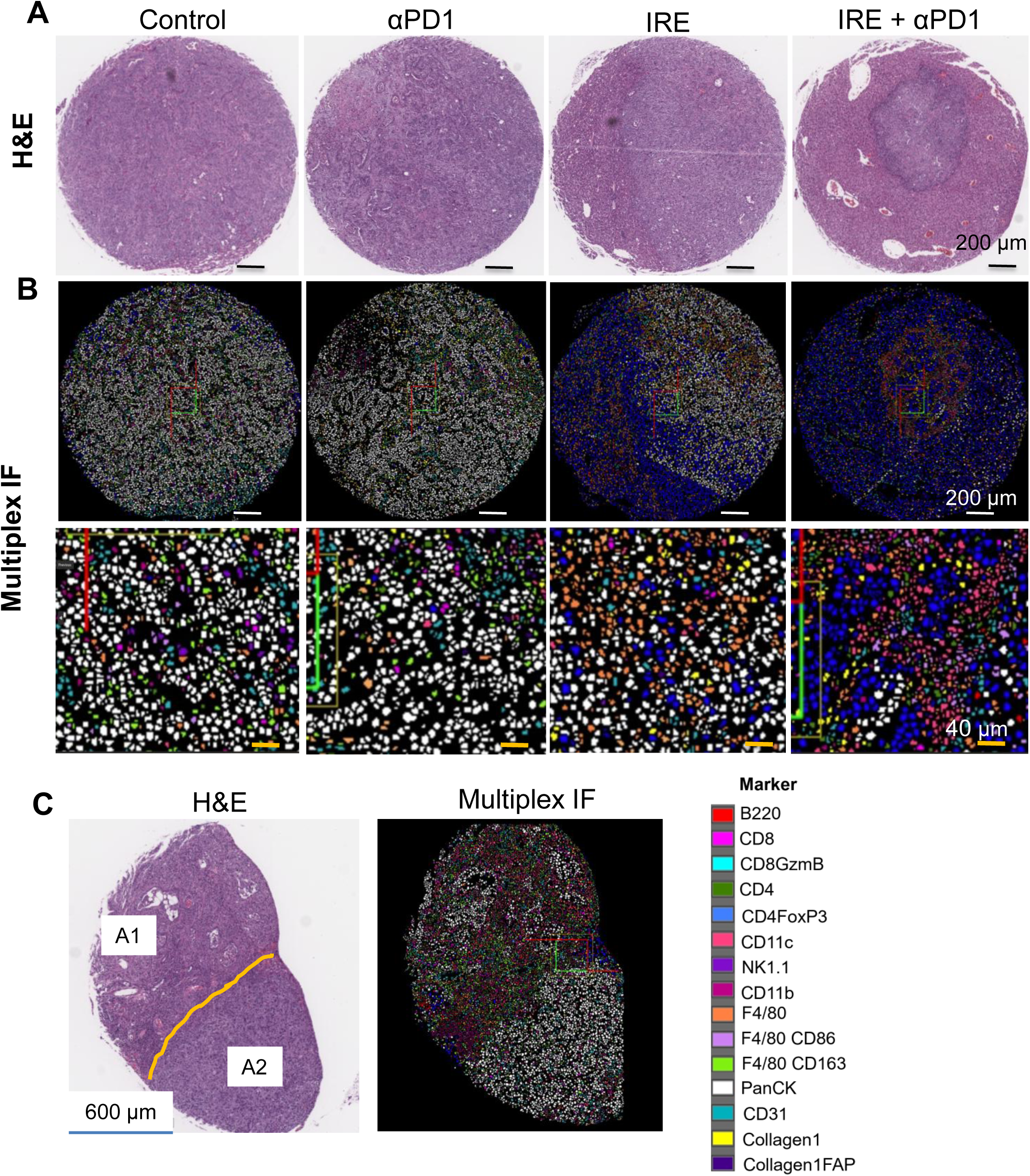
IRE + αPD-1 remodeled the TME of abscopal liver metastases in favor of systemic immune response. **(A, B)** Representative H&E **(A)** and SeqIF images **(B)** of liver tumor TMA slides of each treatment group. **(C)** Representative H&E and SeqIF images of a liver tumor from a mouse treated with IRE alone, showing heterogeneous response after the tumor in the pancreas was treated with IRE. Area 1 (A1) and Area 2 (A2) displayed distinctly different features of immune cell infiltration: A1 was immunologically “hot” with scarce tumor cells, whereas A2 was immunologically “cold” with dense tumor cells. Experimental setup and treatments were the same as outlined in Figure 4A.

In some cases, the response of liver metastases to IRE alone was heterogeneous, as exemplified by the case shown in **Figure 6C**. In this case, there were areas with distinctly different responses. The immunologically “hot” area, area 1 (A1 in **Fig. 6C**), was characterized by abundant immune cells and scarce moderately differentiated tumor area, while the immunologically “cold” area, area 2 (A2 in **Fig. 6C**), was characterized by scarce immune cells and abundant poorly differentiated tumor area. These data indicate that heterogeneous immune cell infiltration into abscopal tumors was a key factor in poor abscopal effects with IRE alone.

The densities of different immune cells in the abscopal liver tumors are shown in **Figure S8**. Similar to what was observed in the primary IRE tumors, densities of immune cells, including F4/80^+^ TAMs, F4/80^+^CD86 M1 TAMs, B220^+^ B cells, CD8^+^ T cells, CD4^+^ T cells, and CD11c^+^ dendritic cells, were significantly higher in the IRE + αPD-1 group than in the IRE and/or αPD-1 monotherapy groups. In contrast, the densities of immunosuppressive F4/80^+^CD163^+^ M2 TAMs and CD4^+^FoxP3^+^ Tregs were significantly lower in the IRE + αPD-1 group. Effective systemic immune response led to depletion of PanCK^+^ tumor cells. Unlike what was observed in the IRE tumors, in the abscopal tumors, there were no differences in the density of CD31^+^ endothelial cells among the different treatment groups, and the densities of αSMA^+^ CAFs and αSMA^+^FAP^+^ CAFs were lower in the IRE-based treatment groups than in the untreated and αPD-1 alone groups (**Fig. S8**).

### scRNA-seq Analysis of Orthotopic KRAS* Tumors Confirmed Upregulation of Immune Response with IRE + αPD-1 versus αPD-1 Monotherapy

In mice with orthotopic KRAS* tumors, tumors were collected and scRNA-seq was performed on day 5 after initiation of treatment with either αPD-1 alone (control) or IRE + αPD-1, and UMAP was performed to analyze cell clusters. The UMAP plot showed that cells were classified into eight major clusters excluding erythroid cells: tumor epithelial cells, T and NK cells, classical dendritic cells, mast cells, myeloid cells, B/plasma cells, fibroblasts, and proliferating cells (**Fig. 7A**). The representative markers for each cell cluster are shown in **Figure 7B**, and populations (i.e., percentages of all cells) of the cell clusters among all cells treated with αPD-1 alone and IRE + αPD-1 are shown in **Figure 7C**. The population of tumor epithelial cells was lower in tumors treated with IRE +αPD-1 (50% of all cells) than in tumors treated with αPD-1 alone (70% of all cells). The total population of immune cells (including T and NK cells, classical dendritic cells, mast cells, myeloid cells, and B/plasma cells) was higher in the IRE + αPD-1 group (40% of all cells) than in the αPD-1 alone group (25% of all cells).

**Figure 7.**
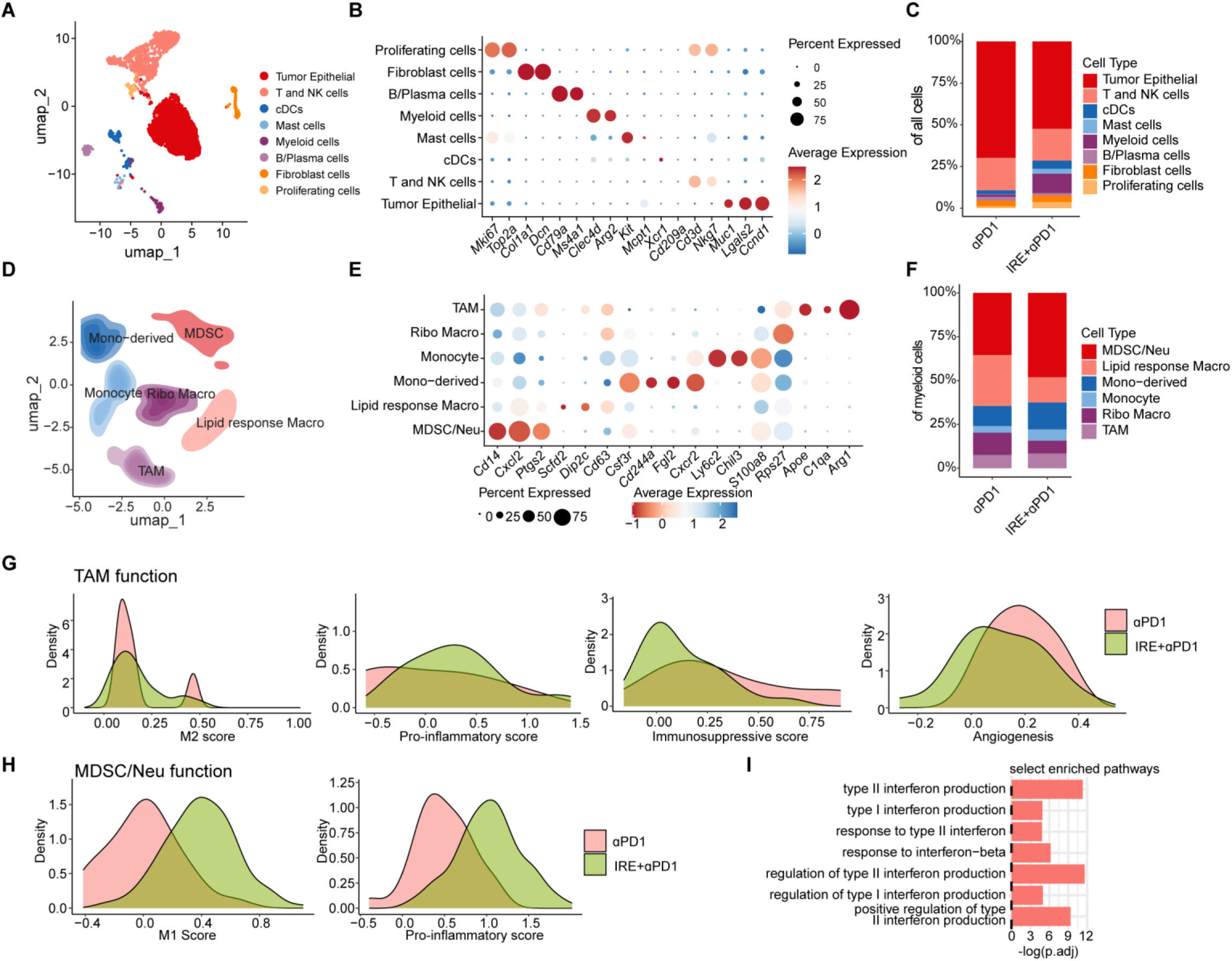
scRNA-seq of orthotopic KRAS* pancreatic tumor on day 5 after initiation of IRE + αPD-1 or αPD-1 alone. **(A)** Uniform Manifold Approximation and Projection (UMAP) plot of major cell clusters in all cells collected from KRAS* tumors. One dot represents one cell. **(B)** Dot plot showing the representative markers for each cluster. The color represents the expression level; the dot size represents the proportion of cells expressing the marker. **(C)** Stacked bar plot showing the populations (i.e., percentages of all cells) of the cell clusters in the αPD-1 alone and IRE + αPD-1 groups. **(D-I)** Results of further analysis of the myeloid cell cluster. **(D)** UMAP plot of subclusters. **(E)** Dot plot showing the representative markers for each cluster. **(F)** Stacked bar plot showing the populations of the myeloid subcluster in the αPD-1 and IRE + αPD-1 groups. **(G)** Distribution plot of TAMs showing the signature scores of M2, pro-inflammatory, immunosuppressive, and angiogenesis functions of TAMs, comparing the IRE + αPD-1 and αPD-1 alone groups. The results showed a subgroup of TAMs in the αPD-1 alone group scored higher in the M2 score. More TAMs in the IRE + αPD-1 group scored higher in pro-inflammatory score and lower in immunosuppressive score, suggesting that IRE + αPD-1 induced a more immunostimulative TAM phenotype than αPD-1 alone. **(H)** Distribution plot of MDSCs showing the signature scores of M1 and pro-inflammatory functions of MDSCs, comparing the IRE + αPD-1 and αPD-1 alone group. **(I)** Bar plots of myeloid cells showing the selected enriched GO pathways of interferon-related processes of genes upregulated in the IRE + αPD-1 group versus the αPD-1 alone group.

Myeloid cells was the cell cluster with the highest difference in population between IRE + αPD-1–treated tumors (11% of all cells) and αPD-1 alone–treated tumors (1% of all cells) (**Fig. 7C**). Analysis of cell subtypes in the myeloid cells cluster revealed that the TAM sub-cluster did not have a significant difference in population between IRE+ αPD-1 (8.4% of all myeloid cells) and αPD-1 alone (7.6% of all myeloid cells) (**Figs. 7D-7F**). Analysis of TAM function showed a lower score of M2 and immunosuppressive TAMs and higher score of pro-inflammatory TAMs with combination therapy compared to αPD-1 monotherapy (**Fig. 7G**). These results suggested that TAMs in their M2 state were reprogrammed into an antitumor M1 state, contributing to an enhanced antitumor effect and abscopal effect of IRE-based combination therapy. These data confirmed the SeqIF imaging data showing that the density of F4/80^+^CD86^+^ M1 TAMs was higher with combination therapy than with αPD-1 alone in both IRE tumors and abscopal tumors.

The percentage of myeloid-derived suppressor cells (MDSCs)/neutrophils was higher in the IRE + αPD-1 group (48.1% of myeloid cells) than in the αPD-1 alone group (35.4% of all myeloid cells) (**Fig. 7F**). Function analysis of the MDSCs/neutrophils cluster showed a higher score of M1-type and pro-inflammatory MDSCs/neutrophils in the combination group than in the αPD-1 alone group (**Fig. 7H**). The enriched GO pathway analysis of myeloid cells showed that interferon-related processes of genes were upregulated in the IRE + αPD-1 group versus the αPD-1 alone group (**Fig. 7I**). Because neutrophils usually demonstrate an M1 score, the higher percentage of MDSCs/neutrophils in the combination group indicated infiltration of neutrophils, which could result from an IRE-induced inflammation response. Moreover, the percentages of lipid response macrophages (lipid-associated macrophages, LAMs) and ribosome biosynthesis–active macrophages (ribo macrophages, RMs) among all myeloid cells were lower in the combination treatment group (14.5% LAMs; 7.3% RMs) than in the αPD-1 alone group (29.1% LAMs; 12.7% RMs) (**Fig. 7F**). LAMs and RMs are in general protumor and immunosuppressive macrophages (31,32). The percentages of cells in the mono-derived and monocyte subclusters among all myeloid cells were higher in the combination treatment group (15.3% mono-derived and 6.5% monocyte) than in the αPD-1 alone group (11.4% mono-derived and 3.8% monocyte) (**Fig. 7F**). Both cell types are usually correlated with antigen-presenting capability to initiate adaptive immune responses. Taken together, these findings indicate that IRE + αPD-1 induced polarization of myeloid cells towards pro-inflammation antitumor phenotypes compared to αPD-1 alone.

T cells and NK cells are crucial players in antitumor immunotherapy, working together in both innate and adaptive immunity to find and destroy cancer cells. The T and NK cell cluster was further analyzed, and subclusters in all T and NK cells are presented in a UMAP plot (**Fig. S9A)** based on the representative markers for each subcluster (**Fig. S9B**). The percentages of each sub cluster in all T and NK cells after αPD-1 alone and IRE + αPD-1 are shown in **Fig. S9C**. Although the total percentage of T and NK cells did not differ between the treatment groups (**Fig. 7C**), the number of proliferating T cells was more than three times as high in the IRE +αPD-1 group (15% of all T and NK cells) as in the αPD-1 alone group (4% of all T and NK cells) (**Figs. S9A-9C**). CD8a^+^ T cells were the main cell population in the proliferating T cell subcluster (**Fig. S9B**).

Subclusters of tumor epithelial cells were analyzed and shown in a UMAP plot (**Fig. S9D**), with the representative markers of each subcluster presented in a dot plot (**Fig. 9E**). The percentages of each subcluster in all tumor epithelial cells after IRE + αPD-1 therapy in comparison with αPD-1 alone are shown in **Figure S9F**. Among these subclusters, the epithelial-mesenchymal transition, tumor-placental growth factor, and tumor-angiogenic subclusters were smaller with IRE + αPD-1 than with αPD-1 alone (**Fig. S9F**). It was previously reported that placental growth factor blockade reduced desmoplasia and tissue stiffness, which resulted in reopening of collapsed tumor vessels and improved blood perfusion while reducing tumor cell invasion (33). Therefore, our findings suggest that combination therapy inhibited the metastasis potential of tumor cells by suppressing epithelial-mesenchymal transition, tumor invasion, and angiogenesis. In addition, the subcluster of tumor cells experiencing stress response was more than twice as large with combination therapy (14.9% of all tumor epithelial cells) as with αPD-1 alone (6.7% of all tumor epithelial cells) (**Fig. S9F**), a finding consistent with *in vitro* RNAseq data (**Table S1**).

Functional enrichment analysis of DEGs of tumor epithelial cells in Gene Ontology - analysis based on biological processes between IRE + αPD-1 (gene ratio > 0) and αPD-1 alone (gene ratio < 0) showed upregulation of the cellular response to interferon-beta and interferon-mediated signaling pathways in tumor epithelial cells after IRE + αPD-1, suggesting that the combination treatment triggered a type I interferon response. Bar plots show the top-ranked GO pathways up-regulated in the IRE + αPD1 group (**Fig. S9G**), as well as the selected interferons-related (**Fig. S9H**) and STING-related (**Fig. S9I**) processes, confirming upregulation of these pathways in IRE + αPD1 combination therapy vs. αPD1 monotherapy.

## DISCUSSION

The abscopal effect induced by localized radiotherapy has been studied extensively (34). However, reports on IRE-induced immune-mediated regression of distant, nonablated tumors are scarce. In this study, we demonstrated that IRE + αPD-1 could induce a strong abscopal effect in preclinical models of PDAC. In the ectopic-ectopic model, IRE + αPD-1 resulted in a durable response in 43% of mice. In the ectopic-orthotopic model, IRE ablation of the subcutaneous tumor in combination with αPD-1 increased median survival compared to each monotherapy, and 20% of the mice treated with IRE + αPD-1 experienced long-term survival. In the clinically relevant orthotopic–liver metastasis model, IRE treatment of tumors in the pancreas in combination with αPD-1 suppressed growth of PDAC tumors in the liver.

Our studies revealed the mechanisms by which IRE-induced local ablation acts as an *in situ* vaccine to induce powerful systemic immune responses. We found that the strong abscopal effects induced by IRE + αPD-1 were driven by IRE-induced immunogenic cell death of PDAC cells, cytosolic release of dsDNA and ssDNA that activated the cGAS-STING pathway, recruitment of myeloid cells and macrophages and their polarization towards M1 phenotype, recruitment of proliferating CD8^+^ T cells, and formation of T cell/B cell clusters (**Fig. 8**).

**Figure 8.**
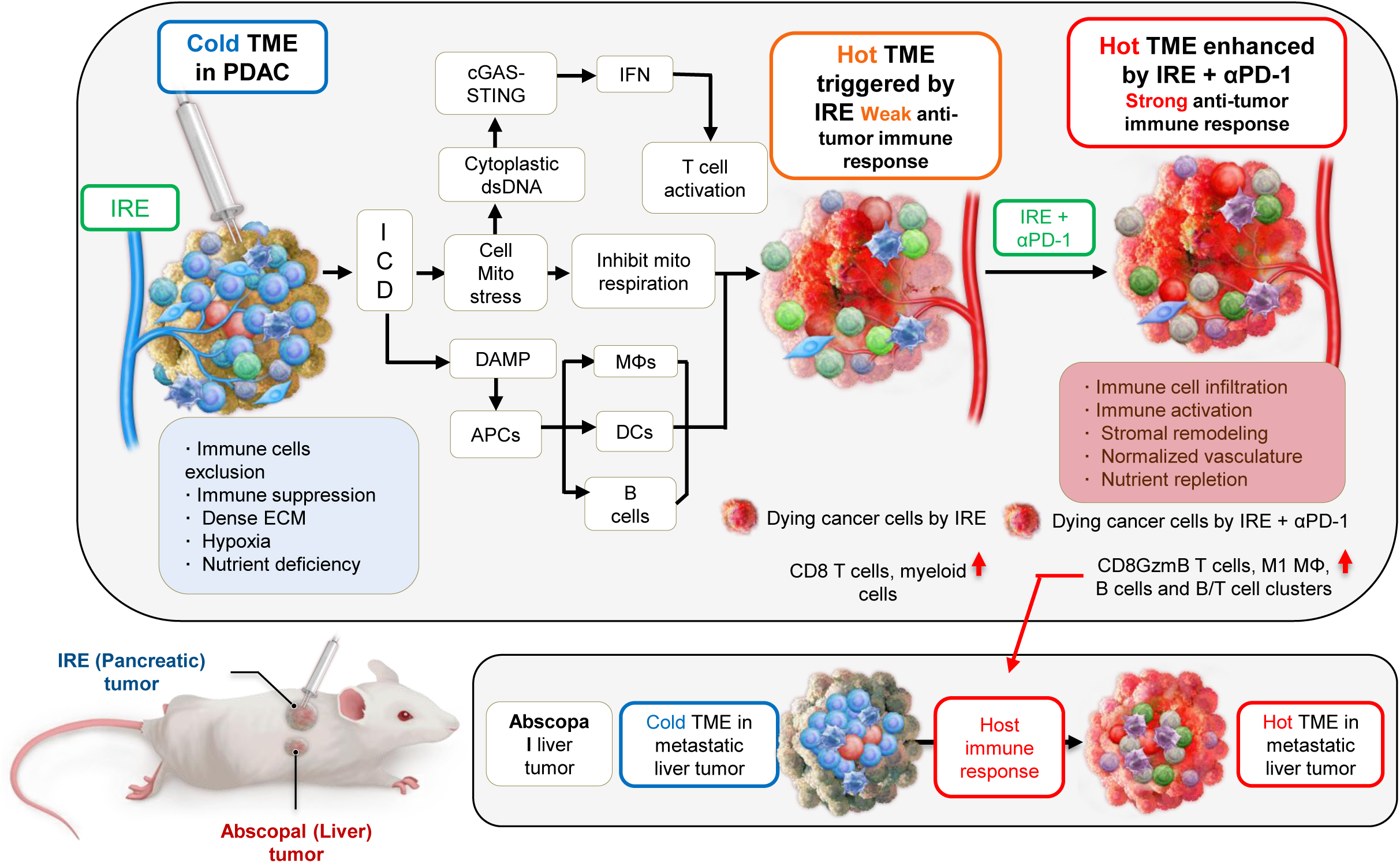
Schematic illustration of mechanisms of action for IRE-induced abscopal effects. APCs, antigen-presenting cells; ICD, immunogenic cell death; IFN, interferon; MΦ, macrophage; mito, mitochondrial.

We first characterized the direct impact of IRE on the transcriptomic changes in KRAS* cells. The top protein-encoding DEGs that were upregulated after IRE were those involved in antigen presentation, stress response, inflammatory immune response and immune cell recruitment, and stroma remodeling. In particular, the gene that encodes Hsp70 was upregulated in both IRE treatment conditions (10p IRE and 20p IRE) and at both timepoints (4 h and 24 h). Hsp70 is recognized as a DAMP “danger signal” and a key component of immunogenic cell death (35). It functions as a molecular chaperone for delivering tumor-derived peptides to antigen-presenting cells to enhance tumor-specific antigen presentation (36). Previously, we showed that IRE caused release of DAMPs and increased the ATP and HMGB1 levels by 11-fold to 13-fold compared to radiotherapy. In a preclinical orthotopic PDAC model, although both 10-Gy radiotherapy + αPD-1 and IRE + αPD-1 prolonged the median survival of mice to approximately 30 days compared to 8 days for αPD-1 monotherapy, no mice were alive at 120 days with radiotherapy + αPD-1, whereas 36% of mice were alive at 120 days with IRE + αPD-1 (6). On the basis of those findings and the findings from our current study, we conclude that IRE played a vital role in the combination therapy in causing rapid onset of immunogenic cell death that releases DAMPs and tumor-associated antigens to activate the adaptive immune system.

GSEA and GO analyses of the RNA-seq data of IRE-treated KRAS* cells showed upregulation of the TNF-α via NF-κB and IL-6/JAK/STAT3 signaling pathways and downregulation of the OXPHOS pathway. To further examine IRE’s role in tumor cell–intrinsic mechanisms of immune response, we assessed the influence of IRE on metabolic activity of KRAS* cells. IRE markedly reduced oxygen consumption rate and extracellular acidification rate and depleted mitochondrial ATP in a pulse number–dependent manner, indicating profound impact of IRE on tumor cells’ mitochondrial function and metabolic activity. IRE uses high-voltage electric pulses to create lasting nanoscale pores in the plasma membrane. Therefore, in addition to causing ATP depletion and mitochondrial dysfunction, IRE may lead to mitochondrial DNA leakage into cytoplasm. Indeed, we observed significantly increased levels of cytosolic dsDNA and ssDNA after IRE treatment of PDAC cells, likely released into cytosol from both mitochondria and nuclei. *In vivo*, cytosolic dsDNA and ssDNA were also detected in KRAS* tumors collected on day 1 and day 4 after IRE treatment. It is known that cytosolic dsDNA and ssDNA activate the immune system through the cGAS-STING sensing pathway (24,25). We found that the expression levels of interferon-stimulated genes downstream of the cGAS-STING pathway, including *Irf1*, *Cxcl10*, and *Isg15*, were all upregulated by IRE. These data indicate that IRE-induced downregulation of the OXPHOS pathway could lead to cytosolic accumulation of mitochondrial DNA and subsequent cGAS-STING activation. Activation of the cGAS-STING pathway can lead to production of pro-inflammatory cytokines, which may have been responsible for the observed activation of the TNF-α via NF-κB and IL-6/JAK/STAT3 signaling pathways (37,38). Analysis of the epithelial cell cluster of the scRNA-seq data showed that compared to αPD-1 alone, combination therapy upregulated interferons and cGAS-STING related genes, findings consistent with our *in vitro* data showing tumor cell–intrinsic mechanisms of immune activation.

IRE + αPD-1 produced stronger abscopal effects and a higher rate of durable response in the ectopic model, in which both the IRE tumors and the abscopal tumors were inoculated subcutaneously, than in the orthotopic models in which either the IRE tumors or the abscopal tumors were grown in the pancreas. This observation suggests that the TME played an important role in the induction of abscopal effects. Orthotopic tumors have a prominent immunosuppressive TME with higher tumor heterogeneity than ectopic tumors. To understand how IRE + αPD-1 impacted the TME of IRE-treated KRAS* tumors in the pancreas, we performed a targeted spatial proteomic profiling using SeqIF imaging. Imaging data from the area proximal to the IRE needles (zones P3 and P4) showed profound transformation of the TME from an immunologically “cold” state to a “hot” state characterized by increased tumor infiltration of macrophages, B cells, and T cells as well as active remodeling of the IRE tumor stroma. In comparison, αPD-1 alone did not cause observable changes in the TME. Although IRE alone increased tumor infiltration of immune cells in the IRE tumors, the increase was less pronounced than with IRE + αPD-1, suggesting that the addition of αPD-1 synergistically enhanced IRE-mediated change in the TME.

Increased infiltration of immune cells in the IRE tumors after IRE + αPD-1 was also observed in zones P1 and P2, which were further away from the IRE needles and the necrotic zone created by IRE. Zones P1 and P2 had significantly fewer F4/80^+^ macrophages, CD11b^+^ myeloid cells, and B220^+^ B cells than zones P3 and P4 had. There were also trends toward fewer CD8^+^/CD4^+^ T cells, F4/80^+^CD86^+^ M1-type macrophages, and cytotoxic CD8^+^GzmB^+^ T cells in zones P1 and P2 than in zones P3 and P4 after IRE + αPD-1 therapy (**Fig. S10A**). For example, comparisons of F4/80 stained macrophages in P2 and P4 zones clearly showed higher tumor infiltration of macrophages in the P4 zone than in the P2 zone (**Fig. S10B)**. These data suggest that the immune response after combination therapy in the IRE-treated tumors in the pancreas was spatially heterogeneous, likely reflecting a heterogeneous gradient of chemokines and cytokines released from treated tumor cells.

Similar to the IRE tumors, the abscopal tumors in the liver showed significantly higher infiltration of immune cells (i.e., F4/80^+^ macrophages, F4/80^+^CD86^+^ M1-type macrophages, CD8^+^/CD4^+^ T cells, and B220^+^ B cells) with IRE + αPD-1 than with either of the two monotherapies. Notably, there were significantly more CD11c^+^ dendritic cells in the IRE + αPD-1 group than in the untreated control group or the αPD-1 alone group, suggesting that IRE was needed to facilitate antigen presentation at the distant site. In addition, there were significantly fewer CD4^+^FoxP3^+^ Tregs in the IRE + αPD-1 group than in the untreated control group and fewer F4/80^+^CD163^+^ M2-type macrophages in the IRE + αPD-1 group than in the αPD-1 alone group, further suggesting that IRE + αPD-1 remodeled the TME of distant liver metastases in favor of dampening immune suppression.

We observed two distinct features after IRE + αPD-1 therapy. First, there was significantly increased population of B220^+^ B cells after IRE + αPD-1 therapy compared to the untreated or αPD-1 monotherapy. Similarly, in the abscopal tumors in the liver, B cell populations were significantly higher in the IRE + αPD-1 group than in the other treatment groups. Moreover, we observed B cells clustering with CD8^+^ T cells in zones P3 and P4 of the IRE tumors after IRE + αPD-1 therapy. B cell and T cell clusters have been observed in the human TME, where they have been described as tertiary lymphoid structures. Tertiary lymphoid structures have been shown to promote antitumor immune responses (39,40). The B cells in tertiary lymphoid structures may act as antigen-presenting cells to activate T cells (41). The formation of B cell/CD8^+^ T cell clusters in PDAC patient tumor samples signified a positive immune response and was associated with better prognosis (42,43). Our scRNA-seq data revealed increased B cell and plasma cell cluster. It is possible that B cells could differentiate into plasma cells that produce tumor-specific antibodies to kill tumor cells. Indeed, B cell-deficient mice exhibited a poor response against established orthotopic KRAS* PDAC tumors after IRE + αPD-1 therapy, demonstrating a crucial role B cell played in mediating strong antitumor activity. Therapeutic strategies that enhance tumor recruitment of B cells and maintain B cell activity may further enhance the complete response rate after IRE + αPD-1 therapy.

A second key feature from the SeqIF imaging studies was significantly increased densities of F4/80^+^ macrophages and F4/80^+^CD86^+^ M1-type macrophages after IRE + αPD-1 therapy compared to the densities in the untreated control and monotherapy groups. αPD-1 alone significantly reduced the population of F4/80^+^CD163^+^ M2 macrophages in both zones P1 and P2 and zones P3 and P4 of the IRE tumors in the pancreas but not in the abscopal tumors in the liver, suggesting that the TME played a role in αPD-1–induced macrophage polarization. It has been reported that αPD-1 could enhance the M1 polarization of TAMs (44). Gordon et al. (45) also showed that αPD-1 could reprogram TAMs to enhance their phagocytic activity. We used *in vitro* Transwell and co-culture systems to study cell migration and phagocytic activity of macrophages. Our results showed that IRE-treated KRAS* cells significantly enhanced macrophage migration. Moreover, co-culture assay showed that RAW264.7 macrophages phagocytosed tumor cells more effectively after IRE treatment of KRAS* cells than after αPD-1 treatment. IRE + αPD-1 induced more F4/80^+^CD86^+^ M1 macrophages than either αPD-1 alone or IRE alone in both IRE tumors and abscopal tumors. Moreover, in tumors from the liver, the density of F4/80^+^CD163^+^ M2 macrophages was lower in the IRE + αPD-1 group than in the untreated control or αPD-1 alone group, suggesting a profound role of IRE in polarizing macrophages in the abscopal tumors. These findings were confirmed by our scRNA-seq data, which showed that compared with αPD1 alone, IRE+αPD1 reduced M2-like phenotype and dampened immunosuppressive and angiogenic programs in TAMs while increasing M1-like and pro-inflammatory features. Taken together, IRE + αPD-1 effectively induced a broader shift toward an immune-activating myeloid state.

Various combination strategies in addition to IRE + αPD-1 for the treatment of PDAC have been tested, with mixed results (6,46,47). Because of the prominent presence of B cells and TAMs in both IRE tumors and abscopal tumors after IRE + αPD-1, approaches that promote polarization of B cells and TAMs in favor of enhanced antigen presentation and activation of T cells are warranted. It is unknown whether and when these macrophages would be educated by the residual tumor cells and change their phenotype to an M2 type. Therefore, the timing of administration of additional drugs after IRE + αPD-1 therapy aimed at polarizing B cells and TAMs is an important issue that must be addressed in future studies.

## METHODS

All animal experiments were approved by the Institutional Animal Care and Use Committee of The University of Texas MD Anderson Cancer Center. Monoclonal anti-mouse PD-1 antibody (CD279 clone: J43) was purchased from BioXCell (West Lebanon, NH, USA). The murine PDAC cell line KRAS* with an inducible *Kras*^G12D^ mutation was obtained from Dr. Ronald A. DePinho’s laboratory (MD Anderson) (48). The murine PDAC cell line HY19636 with a *Kras*^G12D^ mutation was provided by Dr. Haoqiang Ying (MD Anderson). These cell lines were validated by short tandem repeat DNA fingerprinting by the MD Anderson Cytogenetics and Cell Authentication Core using the AmpFLSTR Identifier kit according to the manufacturer’s instructions (Applied Biosystems, Thermo Fisher Scientific). The human pancreatic cancer cell line PANC-1 was purchased from American Type Culture Collection. Hoechst 33342, MitoTracker Red CMXRos, PicoGreen, and OliGreen were purchased from Thermo Fisher Scientific.

### Cell Culture

KRAS* cells were maintained in DMEM–F12 medium supplemented with 10% fetal bovine serum (FBS), 1 µg/mL doxycycline, and penicillin-streptomycin. HY19636 cells and PANC-1 cells were maintained in DMEM supplemented with 10% FBS and penicillin-streptomycin. All cells were cultured in a cell culture incubator with 5% CO_2_ at 37°C.

### IRE Procedure

IRE was performed according to reported protocols (49) using the ECM 830 Square Wave Electroporation System (BTX Harvard Apparatus, Holliston, MA).

#### IRE *in vitro*

Cells were trypsinized and washed with phosphate-buffered saline (PBS) and then resuspended in PBS at 2x10^6^ cells/mL. Aliquots of cells (0.8 mL) were transferred into an Electroporation Cuvette Plus (BTX Harvard Apparatus, Holliston, MA) embedded with two aluminum plate electrodes with a 4-mm gap. The cuvette was placed between the metal contacts of a BTX safety stand (model 630B; BTX Harvard Apparatus), and cell suspension was in direct contact with the plate electrodes and subjected to electroporation at room temperature with the following parameters: voltage, 400 V; pulse duration, 100 µs; pulse repetition frequency, 1 Hz; and number of pulses, 10 or 20. IRE-treated cells or untreated control cells (0.8 mL) were added into 9.2 mL of cell culture medium and incubated for 4 h or 24 h for subsequent assays.

#### IRE *in vivo*

When subcutaneous or orthotopic tumors reached approximately 7 to 8 mm in longest diameter, tumor-bearing mice were anesthetized by inhalation of 1% to 2% isoflurane in O_2_. IRE was performed using a 2-Needle Array electrode (45-0168, BTX Harvard Apparatus) with a 5-mm gap. The electrode was inserted into the center of the tumor along the long axis. The electroporation parameters were as follows: voltage, 1200 V; pulse duration, 100 μs; pulse repetition frequency, 1 Hz; number of pulses, 99. For sham control mice, the needle array was inserted into the center of the tumor, but electroporation was not performed.

### Confocal Microscopy

PANC-1 cells were cultured with DMEM supplemented with10% FBS. Untreated control and IRE-treated (400 V; 20 pulses) PANC-1 cells were seeded on six-well plates with glass cover slides at 37°C in a 5% CO_2_ cell culture incubator for 24 h.

Cytosolic single-stranded DNA (ssDNA) and dsDNA staining were performed as described previously (50). Cells were stained with 5 µg/mL Hoechst 33342 and 100 nM MitoTracker Red CMXRos with or without 10 µg/mL PicoGreen (Invitrogen) or 10 µg/mL OliGreen, and cell plates were incubated for 1 h. After washing with FBS-free cell culture medium, glass cover slides were fixed with 4% paraformaldehyde for 15 min at room temperature. After three washes with PBS, glass cover slides were mounted with DAPI mounting medium (Vector Laboratories, Newark, CA). Fixed slides were kept in a cold room and were protected from light before confocal microscopy examination.

Confocal microscopic images were acquired using a FluoView FV1000 confocal laser scanning microscope equipped with a PLAPO 60x oil objective lens (Olympus, Center Valley, PA).

### Seahorse Cell Mitochondrial Stress Analysis

Mitochondrial stress analysis was performed using a Seahorse XF Cell Mito Stress Test kit (Agilent #103015-100) and a Seahorse XFe96 flux analyzer (Agilent, Santa Clara, CA) according to the manufacturer’s protocol. KRAS* cells were collected and washed with PBS two times, and then the cell concentration was adjusted to 2x10^6^ cells/mL in PBS. KRAS* cells (0.8 mL) were transferred into an electroporation cuvette and treated with IRE. Untreated control or IRE-treated cells were seeded on a poly-D-lysine–coated Seahorse XFe96/XF Pro cell culture plate with 3x10^4^ cells in 80 µL of medium per well and incubated for 4 h at 37°C in a 5% CO_2_ cell culture incubator. Before the assay, the culture medium was replaced with Seahorse XF DMEM (Agilent #103575-100) supplemented with 1 mM pyruvate (Agilent #103578-100), 10 mM glucose (Agilent #103577-100), and 2 mM glutamine (Agilent #103579-100). Cells were incubated at 37°C for 45 min in a non-CO_2_ incubator. Pre-hydrated sensor cartridges were loaded with assay compounds, 1 μM oligomycin, and 0.5 μM rotenone/antimycin. These agents were sequentially injected into the wells to measure changes in oxygen consumption rate and extracellular acidification rate in real time. Upon completion of the assay, nuclei were stained and imaged using Cytation 5 (Agilent BioTek, Santa Clara, CA), and cell counts were obtained using XF Cell Imaging and Counting software (Agilent, version 1.1.0.17). The oxygen consumption rate and extracellular acidification rate data were analyzed with Wave software (Agilent, version 2.6.3.5), and data were normalized by cell numbers.

### RNA-seq

Murine KRAS* PDAC cells were treated with IRE at 400 V with different numbers of pulses. Cells were collected at 4 or 24 h after IRE treatment. RNA of cell lines was isolated using a Direct-zol RNA Miniprep Plus kit (R2072, Zymo Research, Irvine, CA). PolyA enrichment library preparation was performed by Novogene (Sacramento, CA) to generate 250-bp-insert to 300-bp-insert cDNA libraries. RNA-seq was performed by Admera Health (South Plainfield, NJ).

RNA-seq reads quality analysis was performed by FastQC (https://github.com/s-andrews/fastqc). FASTQ files were then mapped to the mouse reference genome (GRCm39) using STAR (v2.7.10a) software (51) and gene expression was quantified by RSEM (v1.3.3) software (52). Processed data files and content were tab-delimited text files containing TPM (transcripts per million) values for each sample. Differential gene expression analysis was conducted by DESeq2 v1.40.2 (53). Gene set enrichment analysis (GSEA) was performed using R packages “fgsea”(v1.26.0, https://github.com/alserglab/fgsea) and “msigdbr” (v 7.5.1, https://github.com/igordot/msigdbr). Figures were generated by “ComplexHeatmap” (v2.18.0) (54) and “ggplot2” (v3.4.2) (55) on R software (v4.3.1).

### Quantitative Real-Time PCR

Total RNA was extracted from KRAS* PDAC cells using the RNeasy Mini kit (#74104, Qiagen, Hilden, Germany) according to the manufacturer’s instructions. cDNA was synthesized using iScript Reverse Transcription Supermix (#1708841, Bio-Rad Laboratories, Hercules, CA). To determine mRNA levels of genes of interest, quantitative real-time polymerase chain reaction (PCR) using gene-specific primers and universal SYBR Green Supermix (#1725274, Bio-Rad Laboratories) was performed. Expression of each gene of interest was normalized to the expression of β-Actin (*Actb)*, a housekeeping gene. Each sample was analyzed three times. Primers used were as follows:

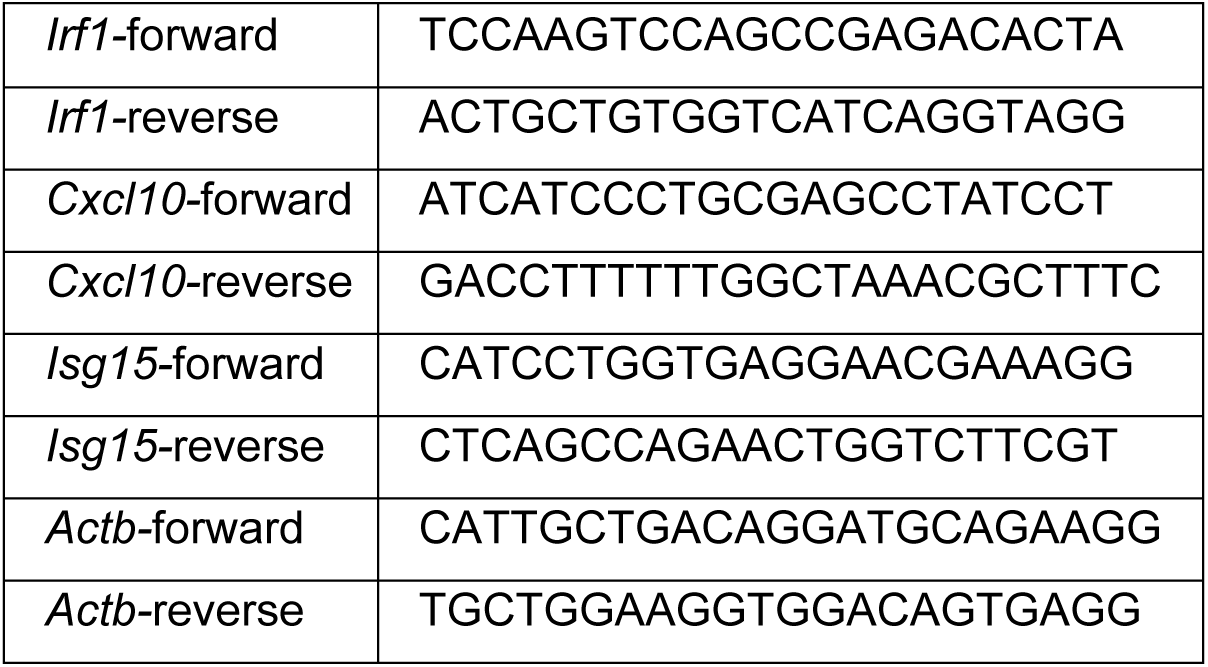

### Animal Model and Treatment

KRAS* PDAC tumor models were established by inoculation of KRAS* cells in female C57BL/6 mice 7 to 8 weeks old (Taconic Biosciences, Germantown, NY). Female B cell deficient Jh C57BL/6 mice (C57BL/6NTac-*Igh-J^em1Tac^*) were a kind gift from Taconic Biosciences.

#### C57BL/6 mice bearing bilateral subcutaneous KRAS* tumors

KRAS* cells were inoculated subcutaneously into the right flank (2x10^6^ cells in 100 μL of PBS) and the left flank (5x10^5^ cells in 50 μL of PBS) at the same time in female C57BL/6 mice. When tumors in the right flank reached approximately 7 to 8 mm in longest diameter, they were treated with IRE with or without αPD-1 injection. Non-IRE-treated mice were used as a control. Growth of right-flank tumors (with or without IRE) and left-flank tumors (untreated abscopal tumors) was monitored by measuring perpendicular diameters of the tumor with a digital caliper until humane euthanasia. The tumor volume was estimated by the formula tumor volume = a × (b^2^)/2, where *a* and *b* are the tumor length and width, respectively, in millimeters. Mice were monitored for survival, with survival defined as the time until the onset of humane endpoint criteria, including a moribund state, hunched posture, labored breathing, ruffled fur, weight loss exceeding 20%, or any tumor larger than 2.0 cm in diameter.

#### C57BL/6 mice bearing subcutaneous and orthotopic KRAS* tumors

KRAS* cells were inoculated subcutaneously into the right flank (2x10^6^ cells in 100 μL of PBS) and orthotopically into the pancreas (1x10^5^ cells in 10 μL of PBS/Matrigel) in female C57BL/6 mice. When the right-flank tumors reached approximately 7 to 8 mm in longest diameter, they were treated with IRE in conjunction with αPD-1 administered intraperitoneally at a dose of 100 μg in 100 μL of PBS at 30 min after IRE and then on days 2, 4, 6, 8, and 10 after IRE for a total of six injections. Mice not treated with IRE or not treated with αPD-1 were used as controls. Tumor growth in the right-flank tumors and the abscopal pancreatic tumors was closely monitored with a digital caliper or bioluminescence imaging until humane euthanasia. Mice were monitored for survival, with survival defined as the time until the onset of humane endpoint criteria, including a moribund state, hunched posture, labored breathing, ruffled fur, weight loss exceeding 20%, or any tumor larger than 2.0 cm in diameter.

#### C57BL/6 mice bearing orthotopic KRAS* tumors and metastases in the liver

KRAS* cells (5x10^5^ cells in 10 μL of PBS/Matrigel) were inoculated into the pancreas to establish an orthotopic tumor, and an aliquot of cancer cells (1x10^5^ cells in 10 μL of PBS/Matrigel) was injected into the right lobe to generate tumor in the liver in female C57BL/6 mice. T2-weighted MRI showed that when KRAS* tumors in the pancreas had reached approximately 7 to 8 mm in longest diameter, the liver tumors were smaller than 1 mm in longest diameter. The primary tumors in the pancreas were treated with IRE in conjunction with intraperitoneal αPD-1 administered intraperitoneally at a dose of 100 μg in 100 μL of PBS at 30 min after IRE and then on days 2, 4, 6, 8, and 10 after IRE for a total of six injections. Mice not treated with IRE or not treated with αPD-1 were used as controls. The liver tumors were not treated with IRE and were used to assess whether there was an abscopal effect. All mice were euthanized 14 days after IRE, and primary tumors in the pancreas and liver tumors in the liver were excised and processed for SeqIF imaging.

### Histological Examination and SeqIF Staining

Tumors from pancreas and liver were harvested and fixed in 10% formalin. Tumor tissue microarrays (TMAs) were prepared by the MD Anderson Veterinary Medicine and Surgery Histology Laboratory. Sequential tissue sections from each TMA blocks are prepared. One slide from each TMA block was stained with hematoxylin and eosin (H&E). SeqIF staining were performed by the Lunaphore COMET system (Lunaphore Technologies SA, Switzerland) on adjacent slides at institutional Flow Cytometry & Cellular Imaging Core. Cellular phenotyping and quantitative analysis were performed with Visiopharm image analysis software (Visiopharm Inc., Broomfield, CO). All antibodies used in the SeqIF staining study are listed in **Table S2**.

### Macrophage Transwell Migration Assay

Migration assays were performed in 6.5-mm Transwell plates with 8-μm pore inserts (product #3422, Corning Inc, Corning, NY) according to the manufacturer’s protocol. After trypsinization and counting, RAW264.7 macrophages (ATCC, Manassas, VA) were resuspended in the medium and added into the upper chamber, and untreated control KRAS* cells or IRE-treated KRAS* cells (400 V, 20 pulses) were added into the lower chamber. Macrophages were allowed to migrate through the insert membrane for 6 h at 37°C under a 5% CO_2_ atmosphere. The inserts were then washed with PBS. Nonmigrating macrophages remaining on the upper surface of the insert were removed with a cotton swab. The migrated cells on the underside of the insert were fixed and stained with crystal violet. The number of migrated macrophages was counted under a light microscope at 40x magnification. The mean number of cells in 10 randomly chosen fields was calculated for each treatment.

### scRNA-seq

C57BL/6 mice were inoculated orthotopically with KRAS* PDAC cells (1x10^5^ cells in 10 μL of PBS/Matrigel). When tumors reached approximately 7 to 8 mm in longest diameter, they were treated with IRE in conjunction with αPD-1 administered intraperitoneally (100 µg/µL/mouse, every other day). Mice treated with αPD-1 alone were used as controls. Pancreatic tumors were collected on day 5 after IRE treatment (after 3 doses of αPD-1).

scRNA-seq was performed as described previously (56). Briefly, tumor tissues were dissociated into single cells with a tumor dissociation Kit (Miltenyi Biotec, Auburn, CA). Cells from three tumors in each treatment group were pooled together for scRNA-seq analysis, which was performed at the MD Anderson Advanced Technology Genomics Core following the default protocol in the user guide of the Chromium Single Cell 5’ Reagent Kit version 1 (10x Genomics, Pleasanton, CA). scRNA-seq libraries were prepared using a Chromium Sing and sequenced using the HiSeq 3000 platform (Illumina Inc, San Diego, CA). Cell Ranger version 3.0.0 (10x Genomics) was used with default parameters to process the reference genome alignment and quantify cells and transcripts. The raw sequencing reads were mapped to the mouse genome.

Clustering and dimension reduction were performed on the filtered Seurat object, which was normalized and scaled with the mitochondrial gene percentage regressed out. The top 2000 most variable genes were identified using the "FindVariableFeatures" function and used for principal component analysis (PCA). Nearest neighbors were identified using the "FindNeighbors" function, and graph-based clustering was performed with "FindCluster" to define cell subtypes. The uniform manifold approximation and projection (UMAP) algorithm was applied for cell subtype visualization. To correct batch effects, the harmony algorithm (R package Harmony) was applied before clustering. Cells were first partitioned into broad categories then further subclustered into subtypes for each category. Cluster identities were assigned based on marker gene expression and manually reviewed to ensure accurate cell type annotation. DEGs among clusters were identified using the "FindAllMarkers" function. The pathway scores were calculated using the gene sets downloaded from the Molecular Signatures Database (MsigDB, https://www.gsea-msigdb.org/gsea/msigdb/). The functional enrichment analysis was conducted by the R package clusterProfier.

### Statistical Analysis

Survival data were analyzed using Kaplan-Meier curves and log-rank tests to compare treatment effectiveness. Student’s t-test or one-way analysis of variance was used to analyze data. Data were expressed as mean ± standard deviation (SD). A p value of less than 0.05 was considered statistically significant. Statistical calculations were performed using GraphPad software (version 10.3.1) and R (version 4.2).

## Supporting Information

**Table S1.**
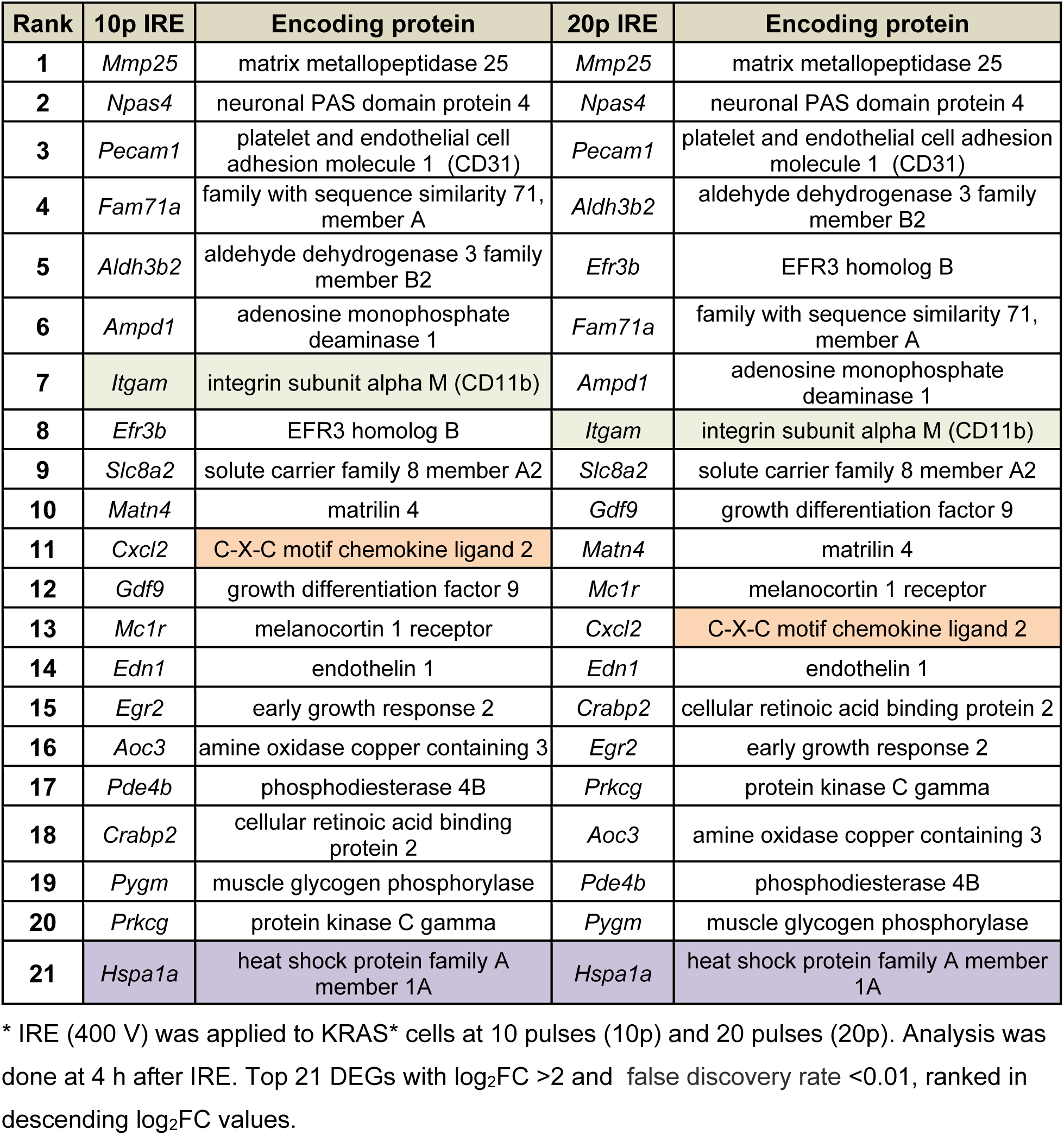
Top 21 upregulated DEGs identified by RNA-seq analysis.

**Table S2.**
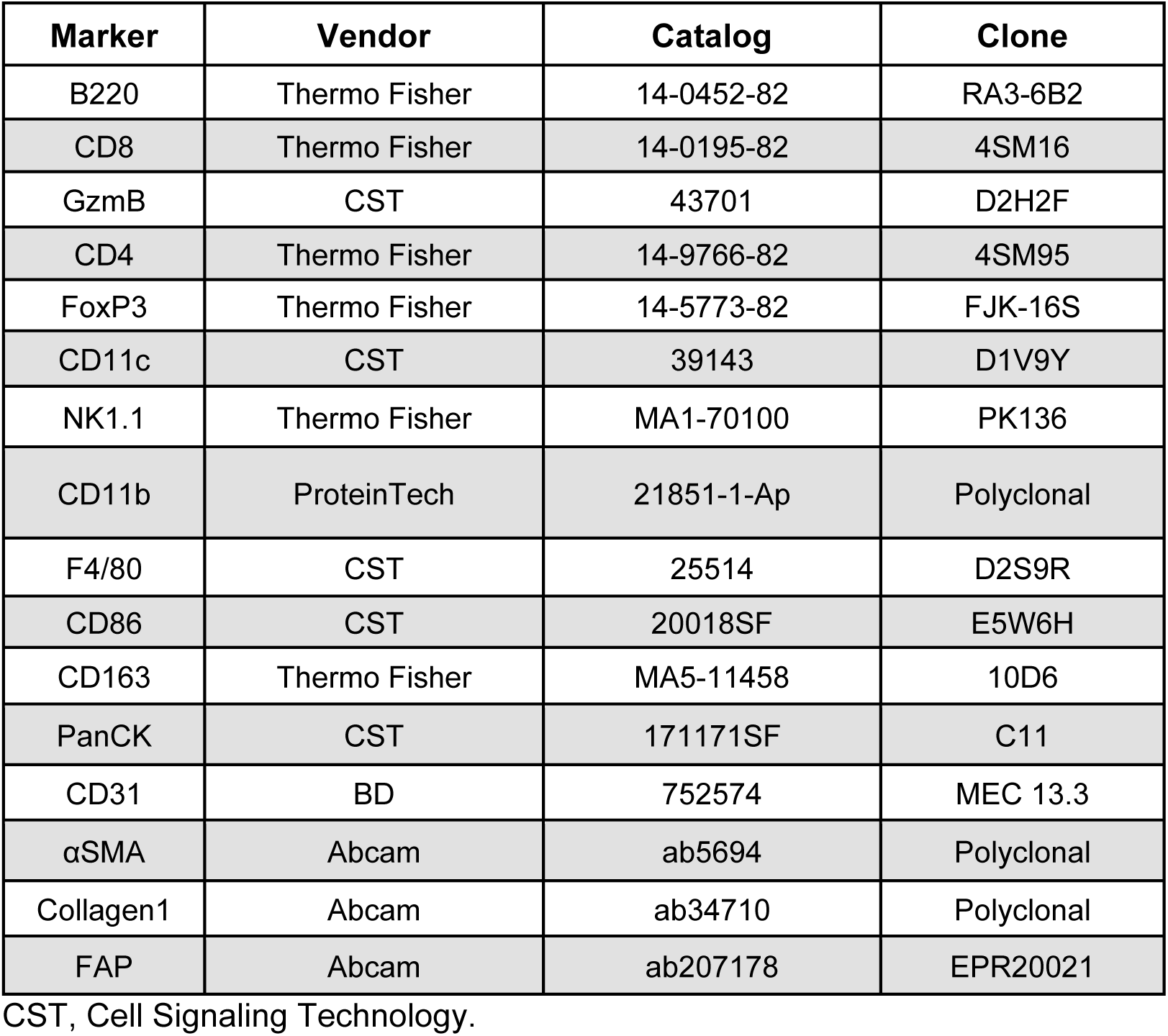
Antibodies used in SeqIF staining.

| **Marker** | **Vendor** | **Catalog** | **Clone** |
| --- | --- | --- | --- |
| B220 | Thermo Fisher | 14-0452-82 | RA3-6B2 |
| CD8 | Thermo Fisher | 14-0195-82 | 4SM16 |
| GzmB | CST | 43701 | D2H2F |
| CD4 | Thermo Fisher | 14-9766-82 | 4SM95 |
| FoxP3 | Thermo Fisher | 14-5773-82 | FJK-16S |
| CD11c | CST | 39143 | D1V9Y |
| NK1.1 | Thermo Fisher | MA1-70100 | PK136 |
| CD11b | ProteinTech | 21851-1-Ap | Polyclonal |
| F4/80 | CST | 25514 | D2S9R |
| CD86 | CST | 20018SF | E5W6H |
| CD163 | Thermo Fisher | MA5-11458 | 10D6 |
| PanCK | CST | 171171SF | C11 |
| CD31 | BD | 752574 | MEC 13.3 |
| αSMA | Abcam | ab5694 | Polyclonal |
| Collagen1 | Abcam | ab34710 | Polyclonal |
| FAP | Abcam | ab207178 | EPR20021 |

CST, Cell Signaling Technology.

## Supporting Information Figure Legends

**Figure S1.** Tumor growth curves for IRE tumors in mice inoculated with KRAS* tumors in both flanks subcutaneously (n=6-7). CR, complete response.

**Figure S2. RNA-seq analysis of KRAS* cells at 4 h and 24 h after IRE treatment.** (**A, B**) Principal component analysis showing gene expression profiles for KRAS* cells at 4 h (**A**) and 24 h (**B**) after IRE (400 V). The plot demonstrates a clear separation between the three experimental conditions based on their gene expression similarities. Control, no treatment; 10p, IRE with 10 pulses; 20p, IRE with 20 pulses. (**C**) Venn diagram showing the numbers of DEGs with |log_2_FC| > 2.0 and false discovery rate < 0.01 in comparisons of different sets of data.

**Figure S3. IRE induced mitochondrial dysfunction.** Effect of IRE on basal respiration (**A**), maximal respiration (**B**), and spare respiratory capacity (**C**). KRAS* cells were analyzed at 4 h after 10p IRE or 20p IRE (400 V). Data are normalized to cell number at the end of the assay (pmol/min/1000 cells) and are expressed as mean ± SD (n = 6/group). OCR, oxygen consumption rate; ECAR, extracellular acidification rate. *p<0.05, **p<0.01, ***p<0.001, ****p<0.0001 (one-way ANOVA).

**Figure S4. IRE induced cytosolic release of ssDNA.** (**A**) Representative confocal fluorescence microscopic images of cytoplasmic ssDNA and corresponding quantification of fluorescence (FL) signal intensity in cytosol and nuclei. PANC-1 cells were stained with OliGreen for ssDNA 24 h after IRE (400 V, 20 pulses). Untreated cells were used as a control. Data are expressed as mean ± SD (n=4). *p<0.05 (Student’s t test). (**B**) Analysis of ssDNA from IRE-treated KRAS* tumors using the Qubit ssDNA assay kit. KRAS* tumor tissues were collected on day 1 and day 4 after IRE (1200 V, 99 pulses) and processed for ssDNA assays. *p<0.05, **p<0.01 (Student’s t test).

**Figure S5. IRE + αPD-1 remodeled the TME to turn immunologically "cold" orthotopic KRAS* PDAC tumors into "hot" tumors.** (**A**) Heatmap of proteomic biomarkers from P3 and P4 spots of TMA slides after indicated treatments. Data are expressed as log_2_(Exp+1). (**B**) Quantitative analysis of cellular biomarkers from SeqIF images. Cell densities are expressed as mean ± SD (n=3-5/treatment x 4 spots/tumor). *p<0.05, **p<0.01 (two-sample Wilcoxon test). Experimental setup and treatment conditions were the same as in Figure 4A.

**Figure S6. IRE-treated KRAS* cells enhanced macrophage migration and phagocytic activity.** (**A, B**) Microphotographs of RAW264.7 macrophages in the upper chamber of a Transwell plate. Cells were co-cultured with untreated tumor cells (**A**) or IRE-treated tumor cells (400 V, 20 pulses) (**B**) seeded in the lower chamber of the Transwell plate. Images were taken at 6 h after co-culture. (**C**) Quantitative analysis of number of migrated macrophages per field of view. Data are presented as mean ± SD (n=18-21). ** p<0.01 (Student’s t test). (**D**) Flow cytometry dot plots of RAW264.7 macrophages co-cultured with IRE-untreated or IRE-treated (400 V, 10 pulses) murine HY19636 PDAC cells. IRE enhanced the phagocytotic activity of macrophages, as shown by higher number of cells in the dual-positive quadrant indicative of F4/80-stained macrophages and CFSE-labeled tumor cells.

**Figure S7. SeqIF images of zones P1 and P2 of KRAS* tumors from pancreas and corresponding data analysis.** (**A, B**) Representative H&E-stained TMA slides (**A**) and corresponding SeqIF images (**B**) of zones P1 and P2 of tumors from each of four treatment groups. (**C**) Quantification of different phenotypes of cells in zones P1 and P2. Cell densities are expressed as mean ± SD (n=3-5 tumors/group x 4 spots/tumor). *p<0.05, **p<0.01, ***p<0.001 (two-sample Wilcoxon test). Experimental setup and treatment conditions were the same as in Figure 4A.

**Figure S8. IRE + αPD-1 significantly impacted the TME of abscopal liver metastases.** Quantification of different immune cells in the TME. Cell densities (cells/mm^2^) are expressed as mean ± SD (n=3-5 tumors/group × 4 spots/tumor). *p<0.05, **p<0.01, ***p<0.001, ****p<0.0001 (two-sample Wilcoxon test). Experimental setup and treatment conditions were the same as in Figure 4A.

**Figure S9. scRNA-seq of orthotopic KRAS* pancreatic tumors on day 5 after initiation of IRE + αPD-1 or αPD-1 alone.** (**A-C**) Results of further analysis of the T/NK cell cluster. (**A**) UMAP plot of subclusters. (**B**) Dot plot showing the representative markers for each cluster. The color represents the expression level; the dot size represents the proportion of cells expressing the marker. (**C**) Stacked bar plot showing the populations (i.e., percentages of all cells) of the T/NK subcluster in the αPD-1 and IRE + αPD-1 groups. There were more than three times as many proliferating T cells in the IRE + αPD-1 group as in the αPD-1 alone group, and most proliferating T cells were CD8a^+^ T cells. (**D-I**) Results of further analysis of the tumor epithelial cell cluster. (**D**) UMAP plot of subclusters. (**E**) Dot plot showing the representative markers for each cluster. (**F**) Stacked bar plot showing the populations (i.e., percentages of all cells) of the tumor epithelial subcluster in the αPD-1 and IRE + αPD-1 groups. (**G**) Bar plots showing the top-ranked functional enrichment of DEGs of tumor epithelial cells in GO biological processes in the IRE + αPD-1 group (gene ratio > 0) versus the αPD-1 alone group (gene ratio < 0). Bar plots showing the selected GO pathways of interferon-related (**H**) and STING-related (**I**) processes of genes upregulated in the IRE + αPD-1 group versus the αPD-1 alone group.

**Figure S10. Analysis of spatial proteomic profiling of selected immune cells in orthotopic KRAS* PDAC tumors treated with IRE + αPD-1.** (**A**) Cell density of selected immune cells comparing Zone P1-P2 versus P3-P4. (**B**) Representative images of tumor-associated macrophages in zone P2 versus zone P4.

## Acknowledgments

We wish to thank MD Anderson Cancer Center colleagues S. P. Deming of the Research Medical Library for editing this manuscript., Thomas Huynh, AS, BS, of the Veterinary Medicine & Surgery Histology Laboratory for assistance with the tissue microarray preparation, Wai-Kin Chan, PhD, of Cancer Biology for help with the Seahorse cell mitochondrial stress analysis, Rony Avritscher, MD, of Interventional Radiology for assistance with liver tumor inoculation, and Kelly Kage, MFA, CMI, of Diagnostic Imaging for her assistance with the artwork. This research was supported in part by grants from the National Cancer Institute (R01CA258540) and the Cancer Prevention and Research Institute of Texas (RP220051). Data were generated in part through the use of the Metabolomics Facility, Research Animal Support Facility, Flow Cytometry and Cellular Imaging Core, Advanced Technology Genomics Core, and Oncology Research and Immuno-monitoring Core, which receive partial support from the National Cancer Institute under grant P30CA016672 to UT MD Anderson Cancer Center. The research reported in this paper was not directly funded through the P30CA016672 grant to UT MD Anderson Cancer Center and is not within the scope of such grant.

## References

1. Mole RH. Whole body irradiation—radiobiology or medicine? Br J Radiol 1953;26:234–41

2. Kingsley DPE. An interesting case of possible abscopal effect in malignant melanoma. Br J Radiol 1975;48:863–6

3. Wersäll PJ, Blomgren H, Pisa P, Lax I, Kälkner K-M, Svedman C. Regression of non-irradiated metastases after extracranial stereotactic radiotherapy in metastatic renal cell carcinoma. Acta Oncol 2006;45:493–7

4. Chang M, Hou Z, Wang M, Li C, Lin J. Recent advances in hyperthermia therapy-based synergistic immunotherapy. Adv Mater 2021;33:e2004788

5. Li D, Shi S, Cao Q, Sood AK, Molldrem JJ, Ma Q, Li C. Copper sulfide nanoparticle-mediated photothermolysis induces immunogenic cell death in ovarian cancer cells. bioRxiv doi: 10.1101/20250708662637 PMID: 40777257; PMCID: PMC12331107 2025

6. Zhao J, Wen X, Tian L, Li T, Xu C, Wen X, et al. Irreversible electroporation reverses resistance to immune checkpoint blockade in pancreatic cancer. Nat Commun 2019;10:899

7. Slone HB, Peters LJ, Milas L. Effect of host immune capability on radiocurability and subsequent transplantability of a murine fibrosarcoma. J Natl Cancer Inst 1979;63:1229–35

8. Demaria S, Ng B, Devitt ML, Babb JS, Kawashima N, Liebes L, Formenti SC. Ionizing radiation inhibition of distant untreated tumors (abscopal effect) is immune mediated. Int J Radiat Oncol Biol Phys 2004;58:862–70

9. Kolberg H-C, Hoffmann O, Baumann R. The abscopal effect: Could a phenomenon described decades ago become key to enhancing the response to immune therapies in breast cancer?. Breast Care (Basel) 2020;15:443–9

10. Totis C, Averbeck NB, Jakob B, Schork M, Volpi G, Hintze DF, et al. Induction of cytoplasmic dsDNA and cGAS-STING immune signaling after exposure of breast cancer vells to X-ray or high-rnergetic varbon ions. Adv Radiat Oncol 2025;10:101783

11. Faraoni EY, O’Brien BJ, Strickland LN, Osborn BK, Mota V, Chaney J, et al. Radiofrequency ablation remodels the tumor microenvironment and promotes neutrophil-mediated abscopal immunomodulation in pancreatic cancer. Cancer Immunol Res 2023;11:4–12

12. Postow MA, Callahan MK, Barker CA, Yamada Y, Yuan J, Kitano S, et al. Immunologic correlates of the abscopal effect in a patient with melanoma. N Engl J Med 2012;366:925–31

13. Twyman-Saint Victor C, Rech AJ, Maity A, Rengan R, Pauken KE, Stelekati E, et al. Radiation and dual checkpoint blockade activate non-redundant immune mechanisms in cancer. Nature 2015;520:373–7

14. Narayanan G. Irreversible electroporation. Semin Intervent Rad 2015;32:349–55

15. Cannon R, Ellis S, Hayes D, Narayanan G, Martin Ii RCG. Safety and early efficacy of irreversible electroporation for hepatic tumors in proximity to vital structures. J Surg Oncol 2013;107:544–9

16. Ringel-Scaia VM, Beitel-White N, Lorenzo MF, Brock RM, Huie KE, Coutermarsh-Ott S, et al. High-frequency irreversible electroporation is an effective tumor ablation strategy that induces immunologic cell death and promotes systemic anti-tumor immunity. EBioMedicine 2019;44:112–25

17. Thomson KR, Cheung W, Ellis SJ, Federman D, Kavnoudias H, Loader-Oliver D, et al. Investigation of the safety of irreversible electroporation in humans. J Vasc Interv Radiol 2011;22:611–21

18. Martin RC, 2nd, Kwon D, Chalikonda S, Sellers M, Kotz E, Scoggins C, et al. Treatment of 200 locally advanced (stage III) pancreatic adenocarcinoma patients with irreversible electroporation: safety and efficacy. Ann Surg 2015;262:486–94; discussion 92-4

19. Udrycka K, Rutkowski K, Osnytskyy A, Małecka-Wojciesko E. Tumor Characteristics and Clinical Features of the Patient as Prognostic Factors in PDAC. Cancers 2025;17

20. Cancer Facts & Figures 2026. American Cancer Society <https://www.cancer.org/content/dam/cancer-org/research/cancer-facts-and-statistics/annual-cancer-facts-and-figures/2026/2026-cancer-facts-and-figures.pdf>.

21. Mannucci A, Goel A. Advances in pancreatic cancer early diagnosis, prevention, and treatment: The past, the present, and the future. CA Cancer J Clin 2026;76:e70035

22. Akhuba L, Tigai Z, Shek D. Major hurdles of immune-checkpoint inhibitors in pancreatic ductal adenocarcinoma. Cancer Drug Resist 2023;6:327–31

23. Panni RZ, Herndon JM, Zuo C, Hegde S, Hogg GD, Knolhoff BL, et al. Agonism of CD11b reprograms innate immunity to sensitize pancreatic cancer to immunotherapies. Sci Transl Med 2019;11

24. Kwon J, Bakhoum SF. The cytosolic DNA-sensing cGAS–STING pathway in cancer. Cancer Discov 2020;10:26–39

25. Yu L, Liu P. Cytosolic DNA sensing by cGAS: Regulation, function, and human diseases. Sig Transduct Target Ther 2021;6:170–15

26. Shim A, Chen Y, Maciejowski J. Activation and regulation of cGAS-STING signaling in cancer cells. Mol Cell 2025;85:3807–22

27. Madkhali OA, Moni SS, Almoshari Y, Sabei FY, Safhi AY. Dual role of CXCL10 in cancer progression: implications for immunotherapy and targeted treatment. Cancer Biol Ther 2025;26:2538962

28. Inzunza J, del Valle AC. Deciphering the liver’s role in pancreatic cancer metastasis: pathways and therapeutic approaches. NPJ Precis Oncol 2025;9:395

29. Hoffmann A, Cheng G, Baltimore D. NF-κB: master regulator of cellular responses in health and disease. Immun Inflamm 2025;1:2

30. Cao Q, Pagel MD, Kingsley CV, Ma J, Long JP, Wen X, et al. Multiparametric MRI to predict response to irreversible electroporation plus anti-PD-1 immunotherapy in pancreatic ductal adenocarcinoma. Magn Reson Med 2026;95:2266–76

31. Marelli G, Morina N, Portale F, Pandini M, Iovino M, Di Conza G, et al. Lipid-loaded macrophages as new therapeutic target in cancer. J Immunother Cancer 2022;10:e004584

32. Metge BJ, Williams La, Swain CA, Hinshaw DC, Elhamamsy AR, Chen D, et al. Ribosomal RNA biosynthesis functionally programs tumor-associated macrophages to support breast cancer progression. Cancer Res 2025;85:1459–78

33. Aoki S, Inoue K, Klein S, Halvorsen S, Chen J, Matsui A, et al. Placental growth factor promotes tumour desmoplasia and treatment resistance in intrahepatic cholangiocarcinoma. Gut 2022;71:185–93

34. Chen X, Yang M, Huang Y, Tu J, Cai Y, Yuan X. Molecular mechanisms underlying the abscopal effect induced by radiotherapy and its synergistic translational potential with immunotherapy. Ther Adv Med Oncol 2025;17:17588359251387534

35. Vaes RDW, Hendriks LEL, Vooijs M, De Ruysscher D. Biomarkers of radiotherapy-induced immunogenic cell death. Cells 2021;10

36. Todryk SM, Gough MJ, Pockley AG. Facets of heat shock protein 70 show immunotherapeutic potential. Immunology 2003;110:1–9

37. Yu L, Liu P. cGAS/STING signalling pathway in senescence and oncogenesis. Semin Cancer Biol 2024;106–107:87-102

38. Chen Q, Sun L, Chen ZJ. Regulation and function of the cGAS-STING pathway of cytosolic DNA sensing. Nat Immunol 2016;17:1142–9

39. Schumacher TN, Thommen DS. Tertiary lymphoid structures in cancer. Science 2022;375:eabf9419

40. Zhao L, Jin S, Wang S, Zhang Z, Wang X, Chen Z, et al. Tertiary lymphoid structures in diseases: immune mechanisms and therapeutic advances. Signal Transduct Target Ther 2024;9:225

41. Rastogi I, Jeon D, Moseman JE, Muralidhar A, Potluri HK, McNeel DG. Role of B cells as antigen presenting cells. Front Immunol 2022;13:954936

42. Helmink BA, Reddy SM, Gao J, Zhang S, Basar R, Thakur R, et al. B cells and tertiary lymphoid structures promote immunotherapy response. Nature (London) 2020;577:549–55

43. Jacquelot N, Tellier J, Nutt Sl, Belz GT. Tertiary lymphoid structures and B lymphocytes in cancer prognosis and response to immunotherapies. Oncoimmunology 2021;10:1900508

44. Dhupkar P, Gordon N, Stewart J, Kleinerman ES. Anti-PD-1 therapy redirects macrophages from an M2 to an M1 phenotype inducing regression of OS lung metastases. Cancer Med 2018;7:2654–64

45. Gordon SR, Maute RL, Dulken BW, Hutter G, George BM, McCracken MN, et al. PD-1 expression by tumour-associated macrophages inhibits phagocytosis and tumour immunity. Nature (London) 2017;545:495–9

46. Long X, Dai A, Huang T, Niu W, Liu L, Xu H, et al. Simultaneous delivery of dual inhibitors of DNA damage repair sensitizes pancreatic cancer response to irreversible electroporation. ACS Nano 2023;17:12915–32

47. Peng H, Shen J, Long X, Zhou X, Zhang J, Xu X, et al. Local Release of TGF-β Inhibitor Modulates Tumor-Associated Neutrophils and Enhances Pancreatic Cancer Response to Combined Irreversible Electroporation and Immunotherapy. Adv Sci (Weinh) 2022;9:e2105240

48. Ying H, Kimmelman Alec C, Lyssiotis Costas A, Hua S, Chu Gerald C, Fletcher-Sananikone E, et al. Oncogenic Kras maintains pancreatic tumors through regulation of anabolic glucose metabolism. Cell 2012;149:656–70

49. Cemazar M, Sersa G, Frey W, Miklavcic D, Teissie J. Recommendations and requirements for reporting on applications of electric pulse delivery for electroporation of biological samples. Bioelectrochemistry 2018;122:69–76

50. Shen YJ, Le Bert N, Chitre AA, Koo CX, Nga XH, Ho SS, et al. Genome-derived cytosolic DNA mediates type I interferon-dependent rejection of B cell lymphoma cells. Cell Rep 2015;11:460–73

51. Dobin A, Davis CA, Schlesinger F, Drenkow J, Zaleski C, Jha S, et al. STAR: ultrafast universal RNA-seq aligner. Bioinformatics 2013;29:15–21

52. Li B, Dewey CN. RSEM: accurate transcript quantification from RNA-Seq data with or without a reference genome. BMC Bioinformatics 2011;12:323

53. Love MI, Huber W, Anders S. Moderated estimation of fold change and dispersion for RNA-seq data with DESeq2. Genome Biol 2014;15:550

54. Gu Z, Eils R, Schlesner M. Complex heatmaps reveal patterns and correlations in multidimensional genomic data. Bioinformatics 2016;32:2847–9

55. Villanueva RAM, Chen ZJ. ggplot2: Elegant Graphics for Data Analysis (2nd ed.). Meas Interdiscip Res Perspect 2019;17:160–7

56. Lin W, Noel P, Borazanci EH, Lee J, Amini A, Han IW, et al. Single-cell transcriptome analysis of tumor and stromal compartments of pancreatic ductal adenocarcinoma primary tumors and metastatic lesions. Genome Med 2020;12:80

