## Supplementary figures for "The abscopal effect of IRE combined with anti–PD-1 achieves local ablation and systemic control of PDAC"

Qizhen Cao et al.

### Supplemental Information

**Table S1. Top 21 upregulated DEGs at 4 h after IRE identified by RNAseq analysis.***

| Rank | 10p IRE | Encoding protein | 20p IRE | Encoding protein |
|---|---|---|---|---|
| 1 | *Mmp25* | matrix metallopeptidase 25 | *Mmp25* | matrix metallopeptidase 25 |
| 2 | *Npas4* | neuronal PAS domain protein 4 | *Npas4* | neuronal PAS domain protein 4 |
| 3 | *Pecam1* | platelet and endothelial cell adhesion molecule 1 (CD31) | *Pecam1* | platelet and endothelial cell adhesion molecule 1 (CD31) |
| 4 | *Fam71a* | family with sequence similarity 71, member A | *Aldh3b2* | aldehyde dehydrogenase 3 family member B2 |
| 5 | *Aldh3b2* | aldehyde dehydrogenase 3 family member B2 | *Efr3b* | EFR3 homolog B |
| 6 | *Ampd1* | adenosine monophosphate deaminase 1 | *Fam71a* | family with sequence similarity 71, member A |
| 7 | *Itgam* | integrin subunit alpha M (CD11b) | *Ampd1* | adenosine monophosphate deaminase 1 |
| 8 | *Efr3b* | EFR3 homolog B | *Itgam* | integrin subunit alpha M (CD11b) |
| 9 | *Slc8a2* | solute carrier family 8 member A2 | *Slc8a2* | solute carrier family 8 member A2 |
| 10 | *Matn4* | matrilin 4 | *Gdf9* | growth differentiation factor 9 |
| 11 | *Cxcl2* | C-X-C motif chemokine ligand 2 | *Matn4* | matrilin 4 |
| 12 | *Gdf9* | growth differentiation factor 9 | *Mc1r* | melanocortin 1 receptor |
| 13 | *Mc1r* | melanocortin 1 receptor | *Cxcl2* | C-X-C motif chemokine ligand 2 |
| 14 | *Edn1* | endothelin 1 | *Edn1* | endothelin 1 |
| 15 | *Egr2* | early growth response 2 | *Crabp2* | cellular retinoic acid binding protein 2 |
| 16 | *Aoc3* | amine oxidase copper containing 3 | *Egr2* | early growth response 2 |
| 17 | *Pde4b* | phosphodiesterase 4B | *Prkcg* | protein kinase C gamma |
| 18 | *Crabp2* | cellular retinoic acid binding protein 2 | *Aoc3* | amine oxidase copper containing 3 |
| 19 | *Pygm* | muscle glycogen phosphorylase | *Pde4b* | phosphodiesterase 4B |
| 20 | *Prkcg* | protein kinase C gamma | *Pygm* | muscle glycogen phosphorylase |
| 21 | *Hspa1a* | heat shock protein family A member 1A | *Hspa1a* | heat shock protein family A member 1A |

**Table S2. Antibodies used in COMET multiplex immunofluorescence staining.**

| Marker | Vendor | Catalog | Clone |
|---|---|---|---|
| **B220** | Thermofisher | 14-0452-82 | RA3-6B2 |
| **CD8** | Thermofisher | 14-0195-82 | 4SM16 |
| **GzmB** | CST | 43701 | D2H2F |
| **CD4** | Thermofisher | 14-9766-82 | 4SM95 |
| **FoxP3** | Thermofisher | 14-5773-82 | FJK-16S |
| **CD11c** | CST | 39143 | D1V9Y |
| **NK1.1** | Thermofisher | MA1-70100 | PK136 |
| **CD11b** | ProteinTech | 21851-1-Ap | Polyclonal |
| **F4/80** | CST | 25514 | D2S9R |
| **CD86** | CST | 20018SF | E5W6H |
| **CD163** | Thermofisher | MA5-11458 | 10D6 |
| **PanCK** | CST | 171171SF | C11 |
| **CD31** | BD | 752574 | MEC 13.3 |
| **αSMA** | Abcam | ab5694 | Polyclonal |
| **Collagen1** | Abcam | ab34710 | Polyclonal |
| **FAP** | Abcam | ab207178 | EPR20021 |

### Figure S1

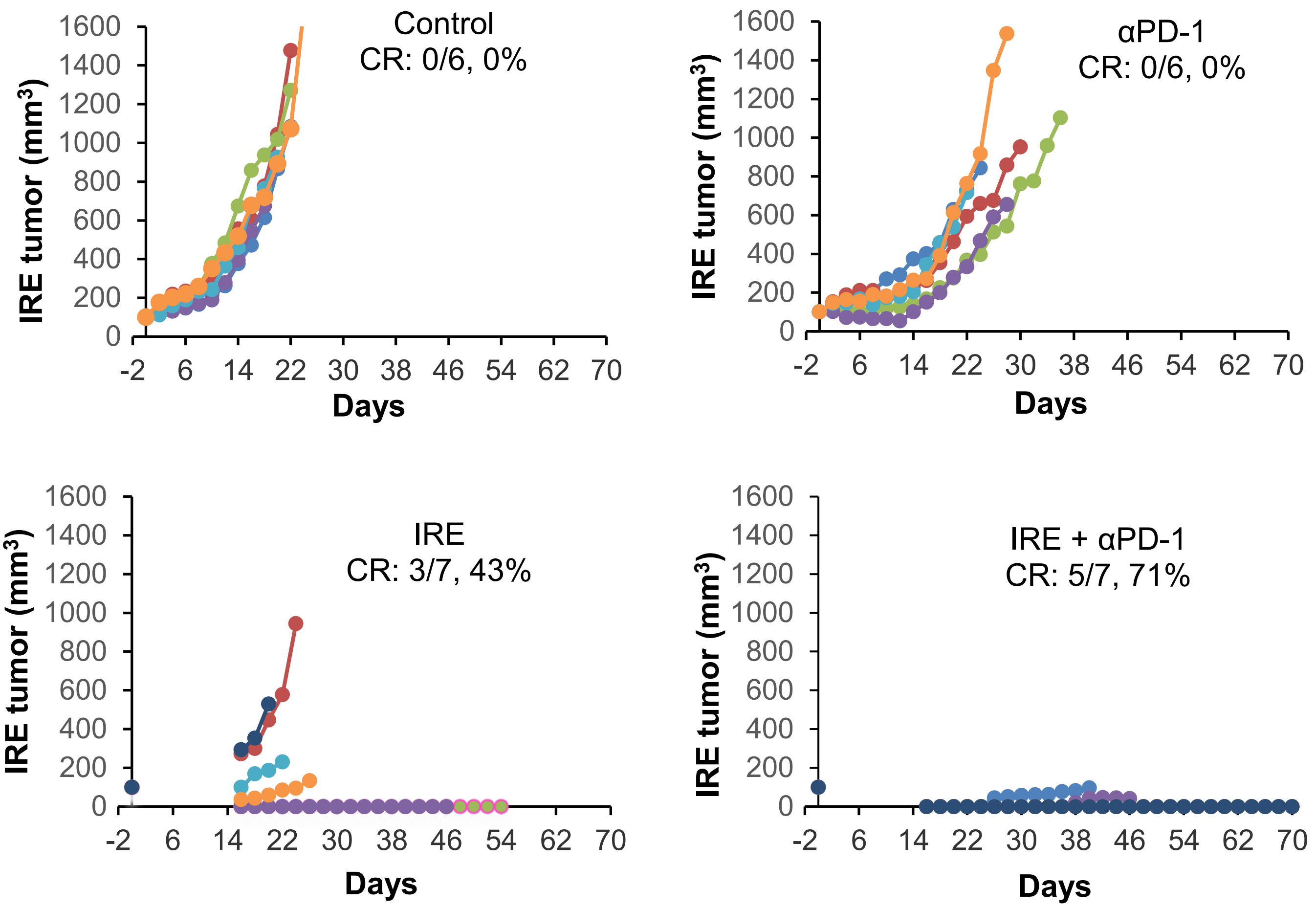


**Figure S1.** Tumor growth curves for IRE tumors in mice inoculated with KRAS* tumors in both flanks subcutaneously (n=6-7). CR, complete response.

**Figure S2**

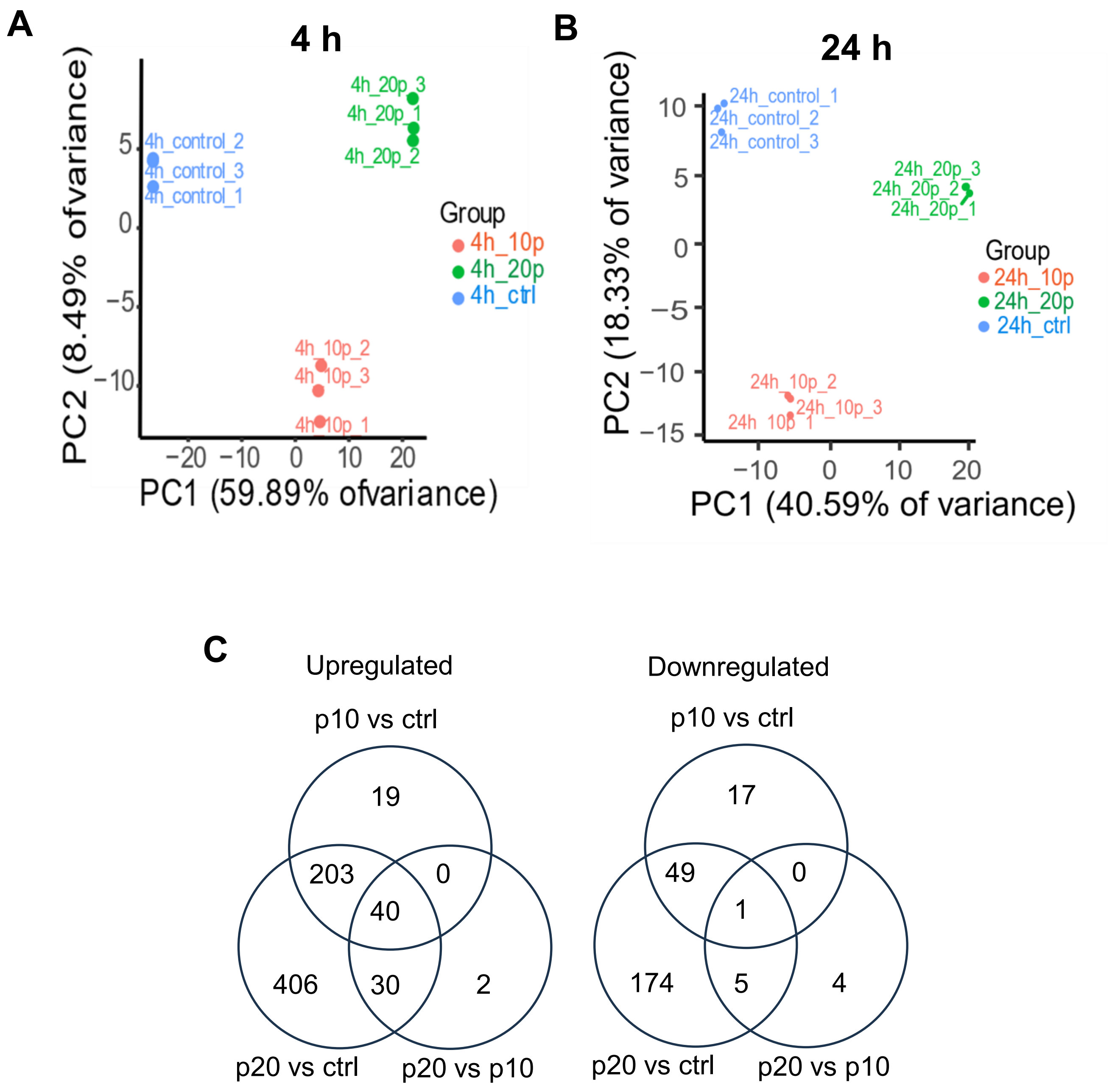


**Figure S2. RNA-seq analysis of KRAS* cells at 4 h and 24 h after IRE treatment.** (**A, B**) Principal component analysis showing gene expression profiles for KRAS* cells at 4 h (**A**) and 24 h (**B**) after IRE (400 V). The plot demonstrates a clear separation between the three experimental conditions based on their gene expression similarities. Control, no treatment; 10p, IRE with 10 pulses; 20p, IRE with 20 pulses. (**C**) Venn diagram showing the numbers of DEGs with $|\log_2 FC| > 2.0$ and false discovery rate < 0.01 in comparisons of different sets of data.

**Figure S3**

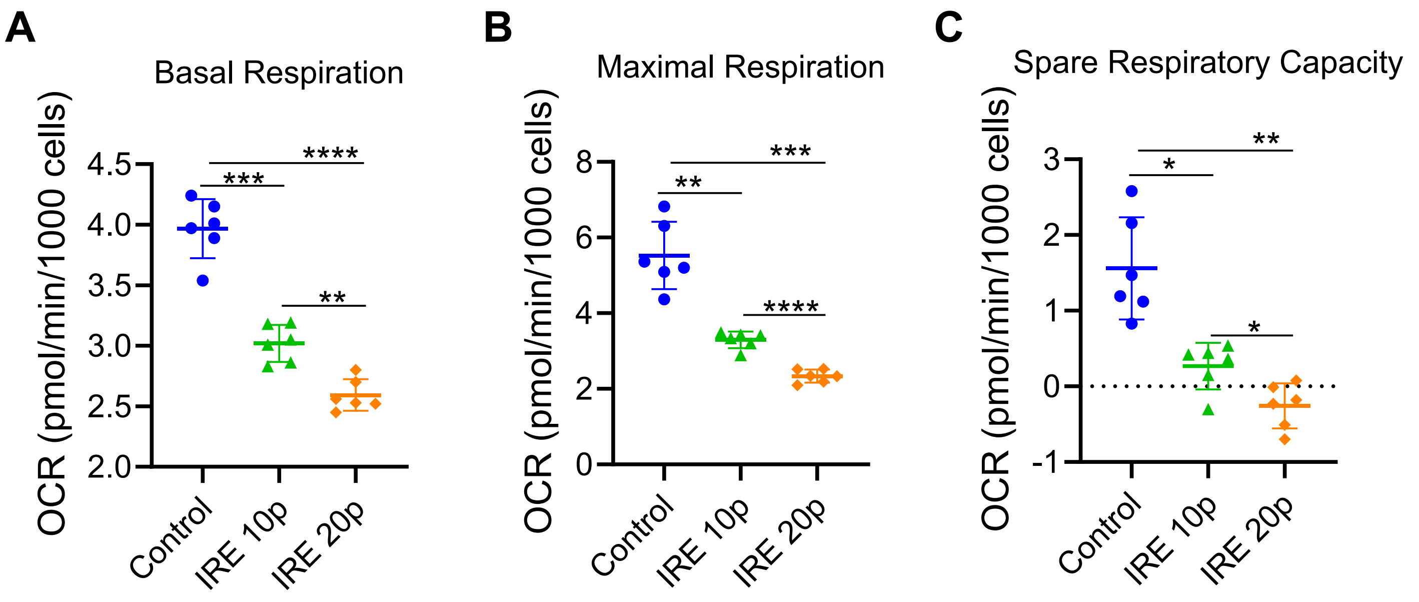


**Figure S3. IRE induced mitochondrial dysfunction.** Effect of IRE on basal respiration (**A**), maximal respiration (**B**), and spare respiratory capacity (**C**). KRAS* cells were analyzed at 4 h after 10p IRE or 20p IRE (400 V). Data are normalized to cell number at the end of the assay (pmol/min/1000 cells) and are expressed as mean ± SD (n = 6/group). OCR, oxygen consumption rate; ECAR, extracellular acidification rate. *p<0.05, **p<0.01, ***p<0.001, ****p<0.0001 (one-way ANOVA).

#### Figure S4

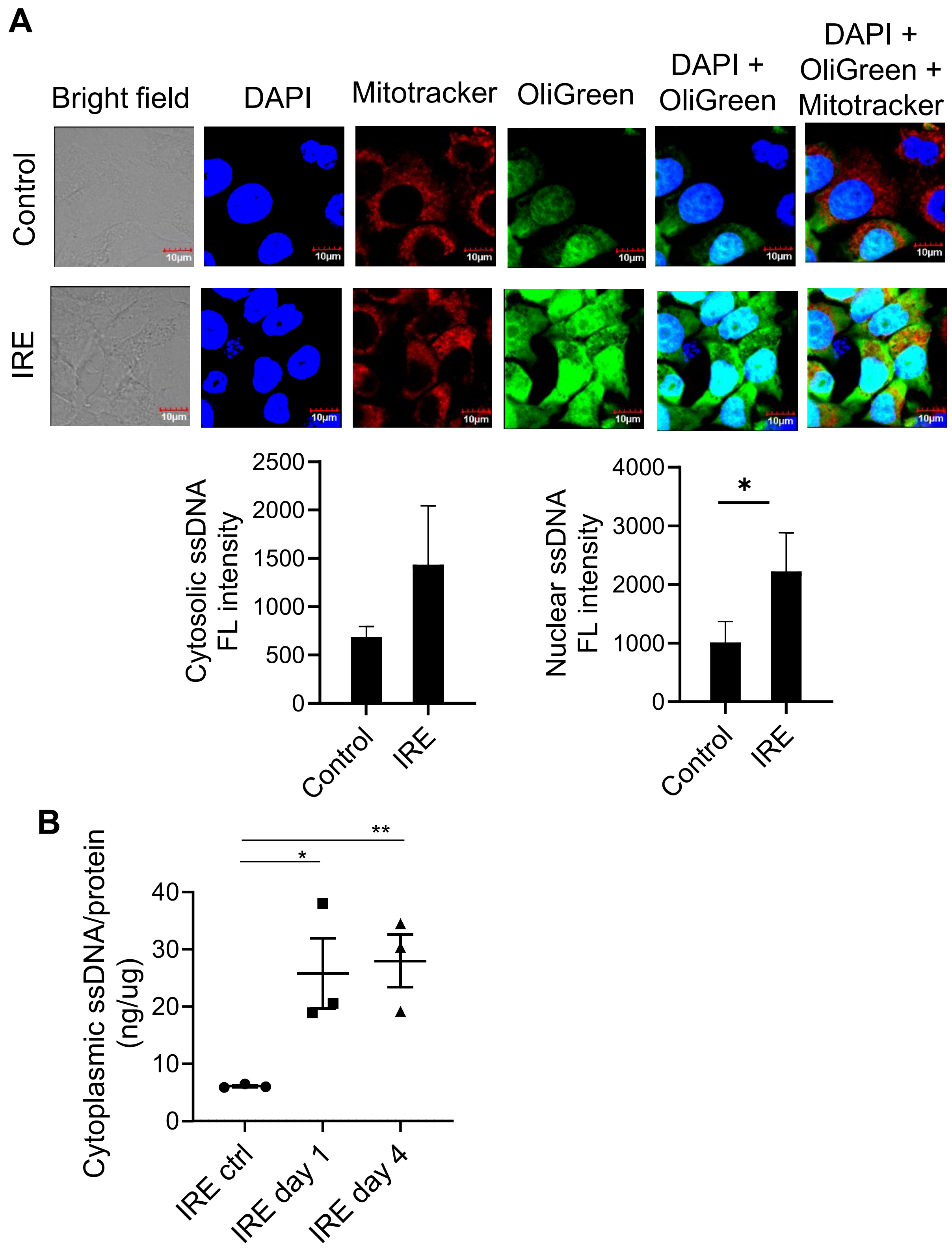


**Figure S4. IRE induced cytosolic release of ssDNA.** (**A**) Representative confocal fluorescence microscopic images of cytoplasmic ssDNA and corresponding quantification of fluorescence (FL) signal intensity in cytosol and nuclei. PANC-1 cells were stained with OliGreen for ssDNA 24 h after IRE (400 V, 20 pulses). Untreated cells were used as a control. Data are expressed as mean ± SD (n=4). *$p<0.05$ (Student's t test). (**B**) Analysis of ssDNA from IRE-treated KRAS* tumors using the Qubit ssDNA assay kit. KRAS* tumor tissues were collected on day 1 and day 4 after IRE (1200 V, 99 pulses) and processed for ssDNA assays. *$p<0.05$, **$p<0.01$ (Student's t test).

### Figure S5

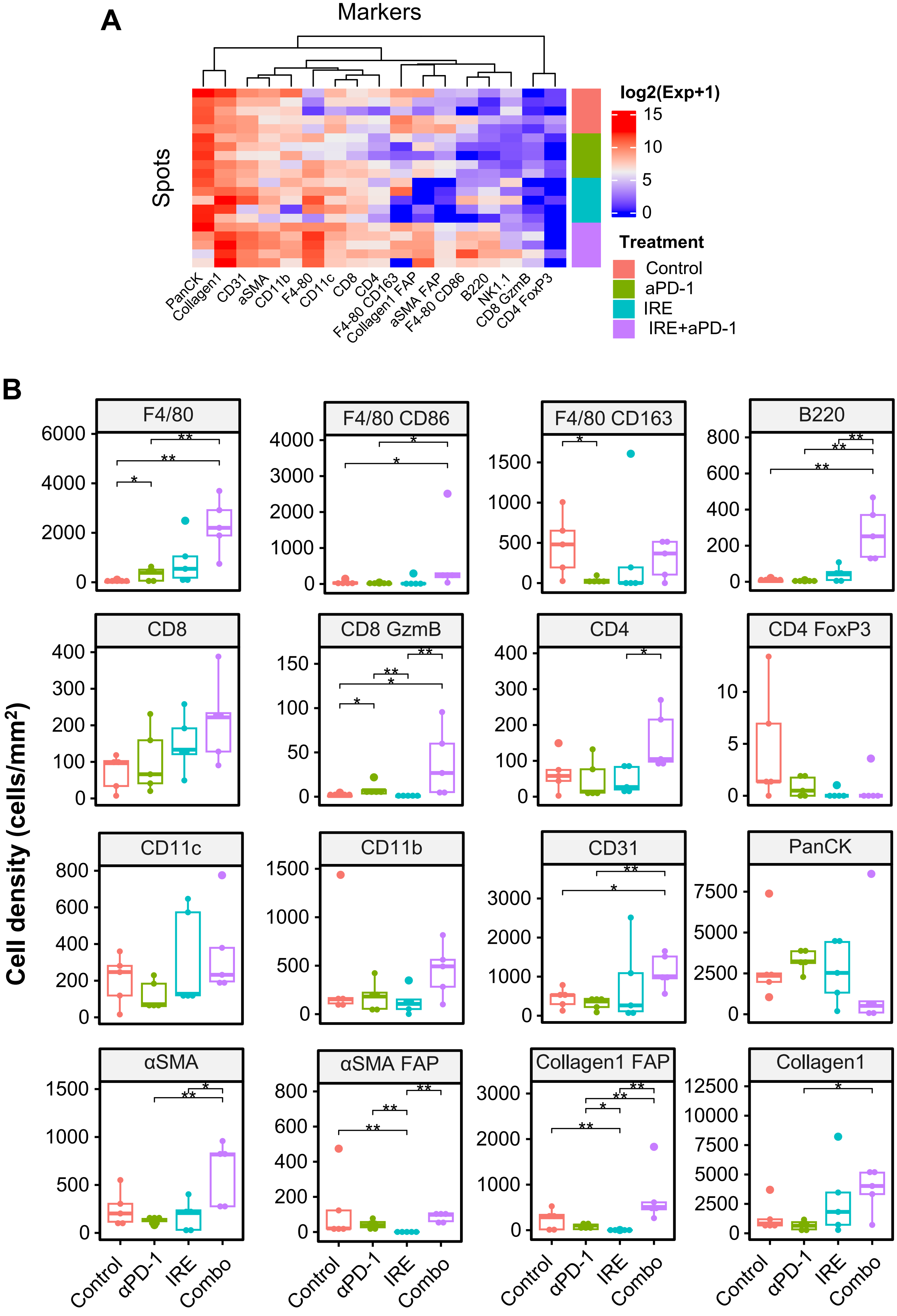
A
Markers
Spots
PanCK
Collagen1
CD31
aSMA
CD11b
F4-80
CD11c
CD8
CD4
F4-80 CD163
Collagen1 FAP
aSMA FAP
F4-80 CD86
B220
NK1.1
CD8 GzmB
CD4 FoxP3
log2(Exp+1)
15
10
5
0
Treatment
Control
aPD-1
IRE
IRE+aPD-1
B
Cell density (cells/mm²)
F4/80
F4/80 CD86
F4/80 CD163
B220
CD8
CD8 GzmB
CD4
CD4 FoxP3
CD11c
CD11b
CD31
PanCK
αSMA
αSMA FAP
Collagen1 FAP
Collagen1
Control
αPD-1
IRE
Combo

**Figure S5. IRE + αPD-1 remodeled the TME to turn immunologically "cold" orthotopic KRAS* PDAC tumors into "hot" tumors.** (**A**) Heatmap of proteomic biomarkers from P3 and P4 spots of TMA slides after indicated treatments. Data are expressed as $\log_2$(Exp+1). (**B**) Quantitative analysis of cellular biomarkers from multiplex IF images. Cell densities are expressed as mean ± SD (n=3-5/treatment x 4 spots/tumor). *$p<0.05$, **$p<0.01$ (two-sample Wilcoxon test). Experimental setup and treatment conditions were the same as in Figure 4A.

### Figure S6

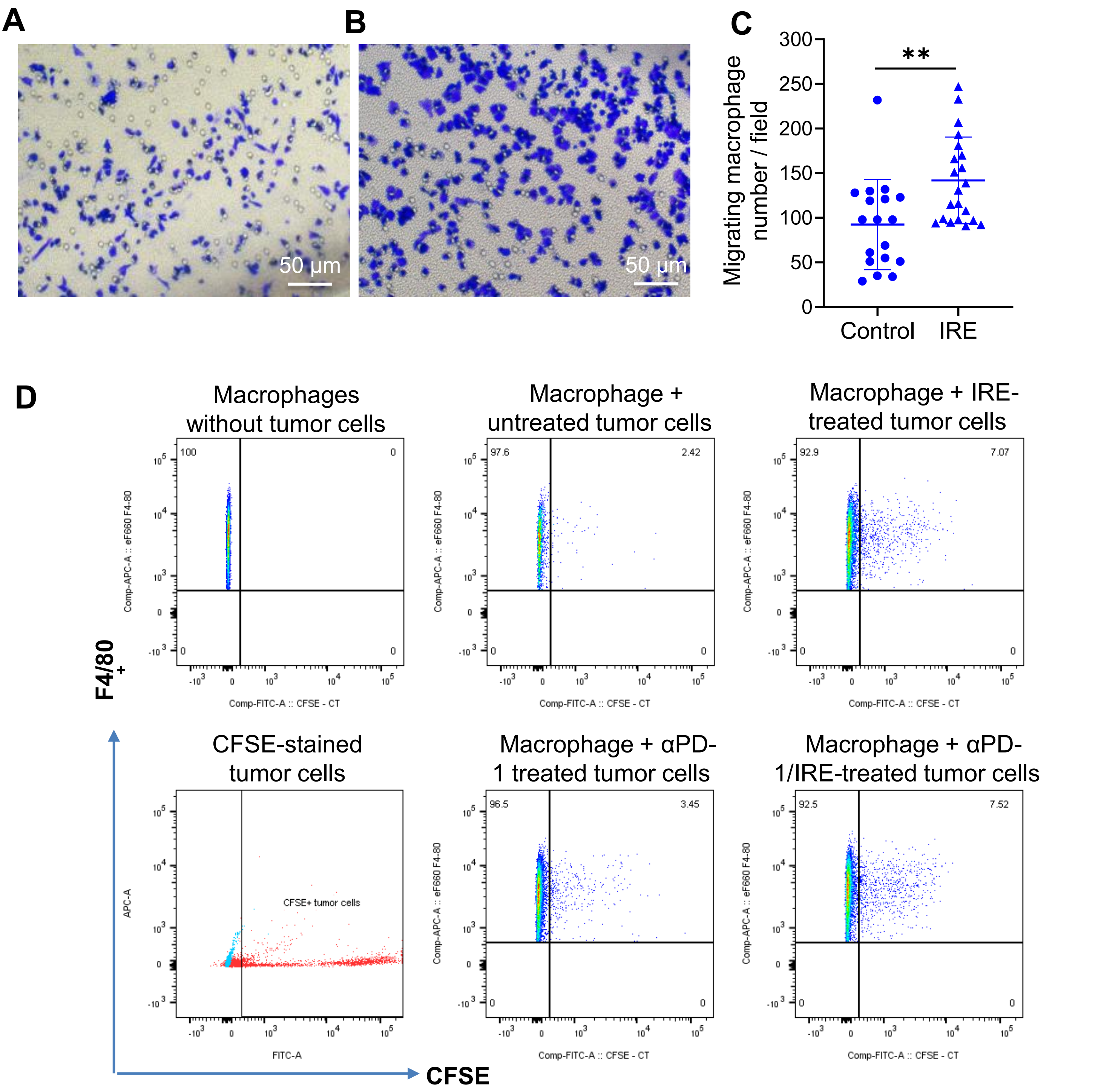


**Figure S6. IRE-treated KRAS* cells enhanced macrophage migration and phagocytic activity.** (**A, B**) Microphotographs of RAW264.7 macrophages in the upper chamber of a Transwell plate. Cells were co-cultured with untreated tumor cells (**A**) or IRE-treated tumor cells (400 V, 20 pulses) (**B**) seeded in the lower chamber of the Transwell plate. Images were taken at 6 h after co-culture. (**C**) Quantitative analysis of number of migrated macrophages per field of view. Data are presented as mean ± SD (n=18-21). ** p<0.01 (Student's t test). (**D**) Flow cytometry dot plots of RAW264.7 macrophages co-cultured with IRE-untreated or IRE-treated (400 V, 10 pulses) murine HY19636 PDAC cells. IRE enhanced the phagocytotic activity of macrophages, as shown by higher number of cells in the dual-positive quadrant indicative of F4/80-stained macrophages and CFSE-labeled tumor cells.

### Figure S7

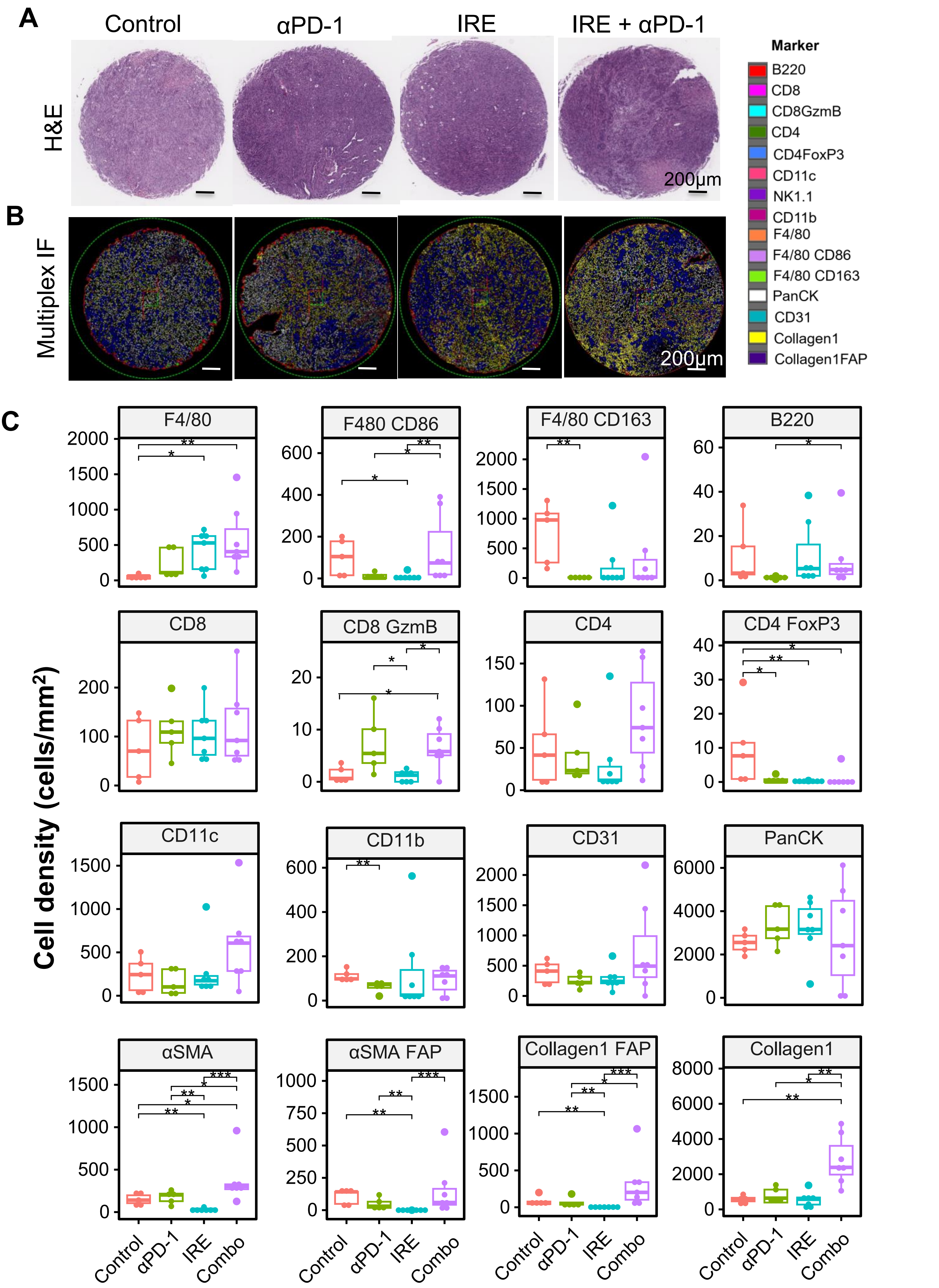

A
Control
αPD-1
IRE
IRE + αPD-1
H&E
200μm
B
Multiplex IF
200μm
Marker
B220
CD8
CD8GzmB
CD4
CD4FoxP3
CD11c
NK1.1
CD11b
F4/80
F4/80 CD86
F4/80 CD163
PanCK
CD31
Collagen1
Collagen1FAP
C
Cell density (cells/mm²)
F4/80
F480 CD86
F4/80 CD163
B220
CD8
CD8 GzmB
CD4
CD4 FoxP3
CD11c
CD11b
CD31
PanCK
αSMA
αSMA FAP
Collagen1 FAP
Collagen1
Control
αPD-1
IRE
Combo

**Figure S7. Multiplex IF images of zones P1 and P2 of KRAS* tumors from pancreas and corresponding data analysis.** (**A, B**) Representative H&E-stained TMA slides (**A**) and corresponding multiplex IF images (**B**) of zones P1 and P2 of tumors from each of four treatment groups. (**C**) Quantification of different phenotypes of cells in zones P1 and P2. Cell densities are expressed as mean ± SD (n=3-5 tumors/group x 4 spots/tumor). *p<0.05, **p<0.01, ***p<0.001 (two-sample Wilcoxon test). Experimental setup and treatment conditions were the same as in Figure 4A.

**Figure S8**

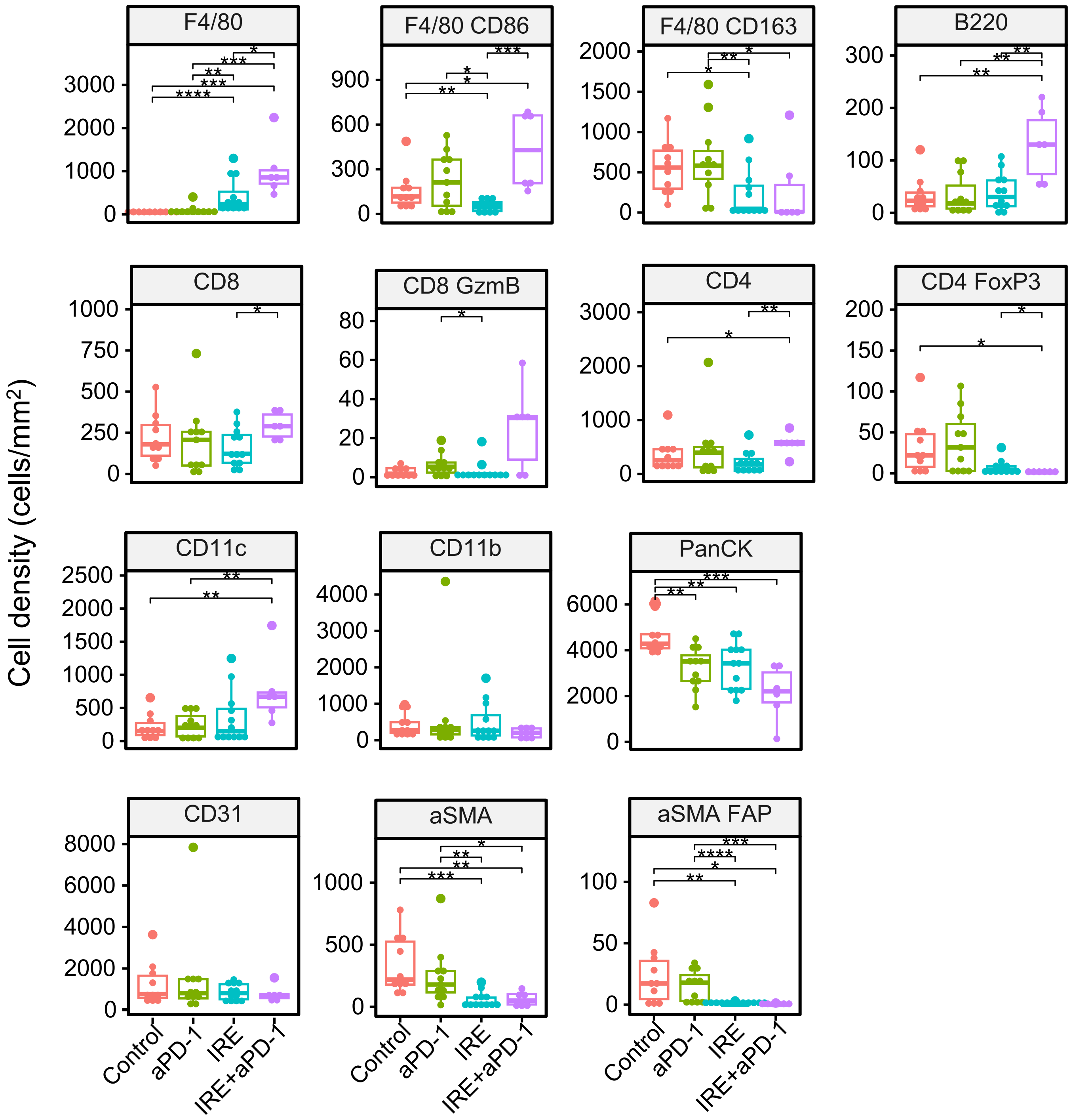


**Figure S8. IRE + αPD-1 significantly impacted the TME of abscopal liver metastases.** (Quantification of different immune cells in the TME. Cell densities (cells/mm$^2$) are expressed as mean ± SD (n=3-5 tumors/group × 4 spots/tumor). *p<0.05, **p<0.01, ***p<0.001, ****p<0.0001 (two-sample Wilcoxon test). Experimental setup and treatment conditions were the same as in Figure 4A.

### Figure S9

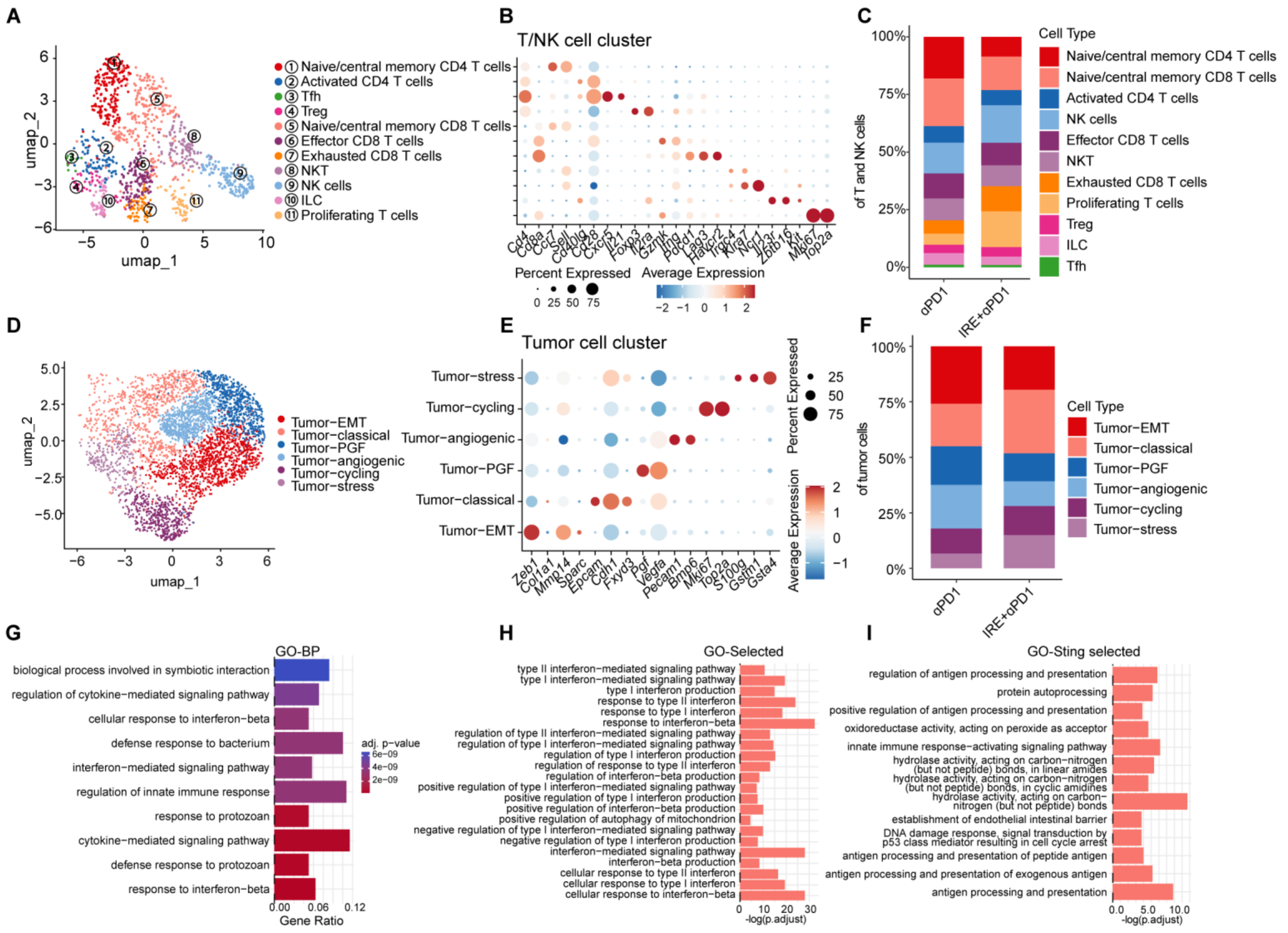


**Figure S9. scRNA-seq of orthotopic KRAS* pancreatic tumors on day 5 after initiation of IRE + αPD-1 or αPD-1 alone.** (**A-C**) Results of further analysis of the T/NK cell cluster. (**A**) UMAP plot of subclusters. (**B**) Dot plot showing the representative markers for each cluster. The color represents the expression level; the dot size represents the proportion of cells expressing the marker. (**C**) Stacked bar plot showing the populations (i.e., percentages of all cells) of the T/NK subcluster in the αPD-1 and IRE + αPD-1 groups. There were more than three times as many proliferating T cells in the IRE + αPD-1 group as in the αPD-1 alone group, and most proliferating T cells were CD8a$^{+}$ T cells. (**D-I**) Results of further analysis of the tumor epithelial cell cluster. (**D**) UMAP plot of subclusters. (**E**) Dot plot showing the representative markers for each cluster. (**F**) Stacked bar plot showing the populations (i.e., percentages of all cells) of the tumor epithelial subcluster in the αPD-1 and IRE + αPD-1 groups. (**G**) Bar plots showing the functional enrichment of DEGs of tumor epithelial cells in GO biological processes in the IRE + αPD-1 group (gene ratio > 0) versus the αPD-1 alone group (gene ratio < 0). Bar plots showing the selected GO pathways of interferon-related (**H**) and STING-related (**I**) processes of genes upregulated in the IRE + αPD-1 group versus the αPD-1 alone group.

### Figure S10

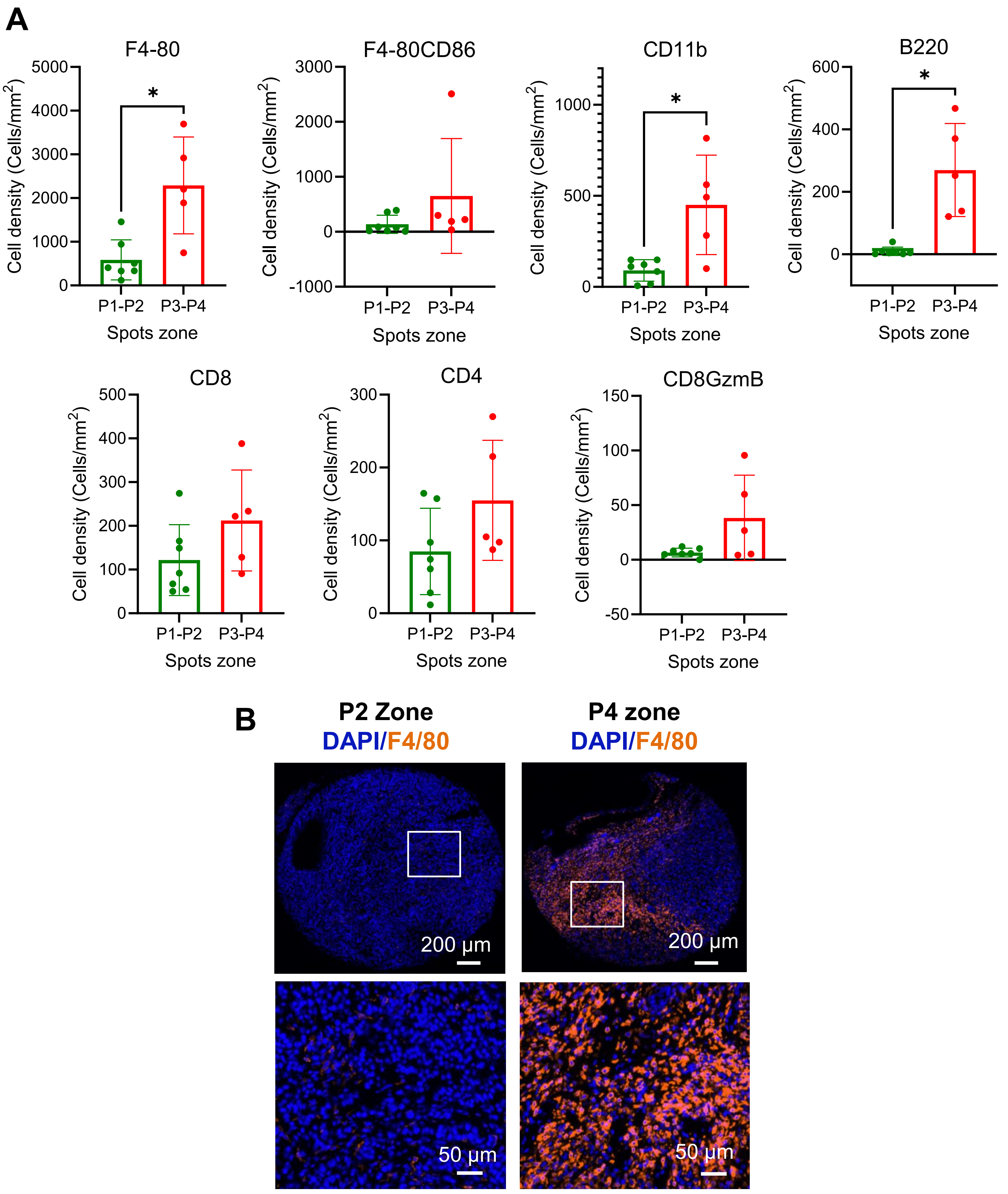


**Figure S10. Analysis of spatial proteomic profiling of selected immune cells in orthotopic KRAS* PDAC tumors treated with IRE + αPD-1.** (A) Cell density of selected immune cells comparing Zone P1-P2 versus P3-P4. (B) Representative images of tumor-associated macrophages in zone P2 versus zone P4.
